# Surface SV2A-Syt1 nanoclusters act as a sequestration hub that limits dynamin-1 recruitment and targeting to recycling synaptic vesicles

**DOI:** 10.1101/2022.12.12.520075

**Authors:** Christopher Small, Callista Harper, Anmin Jiang, Christiana Kontaxi, Nyakuoy Yak, Anusha Malapaka, Elizabeth C. Davenport, Tristan P. Wallis, Rachel S. Gormal, Merja Joensuu, Ramon Martínez-Mármol, Michael A. Cousin, Frédéric A. Meunier

## Abstract

Following exocytosis, the recapture of plasma membrane-stranded vesicular proteins into recycling synaptic vesicles (SVs) is essential for sustaining neurotransmission. Surface clustering of vesicular proteins has been postulated as a ‘pre-assembly’ mechanism for endocytosis – ensuring high-fidelity retrieval. Here, we used single-molecule imaging to examine the nanoclustering of synaptotagmin-1 (Syt1) and synaptic vesicle protein 2A (SV2A) in hippocampal neurons. Syt1 forms surface nanoclusters through interaction of its C2B domain with SV2A, that are sensitive to mutations in this domain (Syt1^K326A/K328A^) and knocking down SV2A. SV2A co-cluster with Syt1 and blocking SV2A’s cognate interaction with Syt1 (SV2A^T84A^) also decreased SV2A clustering. Surprisingly, impairing SV2A-Syt1 nanoclustering enhanced plasma membrane recruitment of key endocytic protein dynamin-1, leading to accelerated Syt1 endocytosis, altered intracellular sorting and decreased trafficking of Syt1 to Rab5-positive endocytic compartments. SV2A-Syt1 surface nanoclusters therefore negatively regulate the rate of their own re-entry into recycling SVs by controlling the recruitment of the endocytic machinery.

## Introduction

Synaptic vesicle (SV) recycling involves a balance between fusion (exocytosis) and retrieval (endocytosis) of SVs from the plasma membrane (PM) at nerve terminals during neurotransmission. Both exocytosis and compensatory endocytosis involve the coordinated actions of proteins and lipids to ensure high fidelity of vesicular protein retrieval during high rates of SV fusion. To sustain neurotransmission, nascent recycling SVs need to recapture essential vesicular machinery that are stranded at the PM. However, the mechanisms through which neurons retrieve essential vesicular proteins from the PM are not well defined. As certain SV proteins lack canonical recognition motifs for endocytic adaptor molecules, interactions between vesicular cargoes may facilitate recruitment of proteins from the PM into SVs, thus preserving vesicle protein stoichiometry (Takamori et al., 2006; Wilhelm et al., 2014) during neurotransmission. For example, vesicle-associated membrane protein 2 (VAMP2) is a soluble N-ethylmaleimide-sensitive factor attachment protein receptor (SNARE) family member that regulates the fusion of SVs with the PM (Südhof and Rothman, 2009), whose internalisation is facilitated in part, via its interaction with synaptophysin (Gordon and Cousin, 2013; Gordon et al., 2011; Harper et al., 2021; Harper et al., 2017). Similarly, vesicular glutamate transporter 1 (vGlut1) facilitates the recruitment of multiple SV proteins from the PM into SVs (Pan et al., 2015). Thus, interactions between vesicular molecules are theorised to improve the fidelity of endocytic uptake and allow SVs to retain their discrete protein content during multiple rounds of fusion.

One mechanism through which protein interactions improve the fidelity of endocytosis is by forming nanoclusters. Following exocytosis, vesicular proteins cluster at the PM through protein-protein interactions. Notably, VAMP2 disperses following exocytosis and subsequently re-clusters via interactions with endocytic proteins, particularly AP180 and clathrin assembly lymphoid myeloid leukemia (CALM) (Gimber et al., 2015). The endocytic machinery therefore has the potential to initiate clustering of surface-stranded vesicular proteins. However, it is not clear what factors control the clustering of other vesicular proteins, such as synaptotagmin-1 (Syt1). Syt1 is a transmembrane SV molecule that is involved in calcium (Ca^2+^)-dependent exocytosis (Geppert et al., 1994) and forms clusters at the PM (Opazo et al., 2010; Willig et al., 2006). Syt1 binds to the PM phospholipid phosphatidylinositol-4,5-bisphosphate in a Ca^2+^-dependent manner through its cytosolic C2A and C2B domains (Bai et al., 2002; Schiavo et al., 1996; Stein et al., 2007) to mediate exocytosis. Syt1 forms a complex with another vesicular transmembrane protein, synaptic vesicle protein 2A (SV2A) (Bennett et al., 1992), which comprises twelve transmembrane-spanning domains capped by cytosolic C-terminal and N-terminal regions. The SV2A-Syt1 interaction occurs via the cytosolic domains of Syt1 and SV2A: hydrogen bonds are formed between two lysine residues (K326/K328) residing in the polybasic region of Syt1’s Ca^2+^-binding C2B domain (Fernandez et al., 2001) and the T84 epitope on the N-terminus of SV2A upon phosphorylation of the T84 epitope by casein kinase 1 family kinases (Zhang et al., 2015). SV2A has an enigmatic function in neurotransmission, which involves controlling Syt1 trafficking in neurons. SV2A interacts with Syt1 following membrane fusion (Wittig et al., 2021) and controls the retrieval of Syt1 during endocytosis (Kaempf et al., 2015; Zhang et al., 2015). Notably, SV2A knockout reduces Syt1 levels in SVs and at the PM (Yao et al., 2010). For these reasons, SV2A is also a strong candidate as a regulator of Syt1 nanoclustering during SV recycling.

In this study, we investigated the role of protein-protein interactions in controlling the surface clustering of vesicular machinery at the PM, and how these interactions and clustering events facilitate the re-entry of proteins into recycling SVs during endocytosis. We hypothesised that nanoclustering of vesicular proteins at the PM allows for the generation of a ‘readily-accessible’ pool of pre-assembled molecules, forming a depot from which vesicular proteins may be selectively retrieved into nascent recycling SVs. Using super-resolution imaging, we identified the determinants of Syt1 and SV2A nanoclustering by manipulating interactions between Syt1 and SV2A, as well as interactions with endocytic machinery. Syt1-SV2A interaction was shown to be critical for the surface nanoclustering of their respective partner, whilst manipulation of the endocytic machinery had no effect on the nanoclustering of either molecule. Blocking SV2A-Syt1 nanoclustering accelerated Syt1 retrieval during SV endocytosis. This manipulation also led to increased mobility of internalised Syt1 suggesting alterations in recycling SV nanoscale organisation. The findings presented in this study suggest that Syt1 is dynamically sequestered into nanoclusters in an activity-dependent manner through its interaction with the N-terminal tail of SV2A, and that the nanoclustering of Syt1 by SV2A decreases the kinetics of Syt1 endocytic reuptake. Accordingly, we report that SV2A interaction also controls the organisation of Syt1 following internalisation, causing Syt1 entrapment within endocytic pathways associated with early/bulk endosome formation. SV2A therefore plays a critical role in the recycling of Syt1, with implications for the function of Syt1 during multiple rounds of vesicle fusion.

## Results

### Interaction of the Syt1 C2B domain (K326/K328) with SV2A controls the activity-dependent confinement of PM-stranded Syt1 in nerve terminals

First, we investigated the surface mobility and nanoclustering of Syt1 in murine primary cultures of hippocampal neurons using universal Point Accumulation Imaging in Nanoscale Topography (uPAINT). This single-particle tracking technique allows for selective analysis of the nanoscale organisation of surface proteins via labelled ligand tracking (Giannone et al., 2010; Giannone et al., 2013; Joensuu et al., 2016). To visualise Syt1 on the surface of hippocampal neurons, we overexpressed Syt1 tagged with pHluorin (Syt1-pH; Fig. 1A i) – a pH-sensitive green fluorescent protein (GFP) that is quenched in the acidic SV environment and unquenched following exocytosis due to exposure to the neutral, extracellular pH (Miesenböck et al., 1998). Epifluorescence imaging of wild-type Syt1 (Syt1^WT^-pH) revealed distinct synaptic boutons that lined the axon of mature neurons (DIV18) (Fig. 1A ii). To track single Syt1-pH molecules at nanoscale level on the PM, we applied atto647N-labelled anti-GFP nanobodies (NBs) (Kubala et al., 2010) in a depolarizing buffer to increase the level of SV fusion with the PM, and therefore the amount of Syt1-pH on the PM available for atto647N-NB binding (Fig. S1). The atto647N fluorophore was excited (647 nm) in total internal reflection fluorescence (TIRF) mode, to selectively image the surface population of atto647N-NB-bound Syt1^WT^-pH (16,000 frames, 320 s) (Fig. 1B). Within nerve terminals, we observed the presence of distinct Syt1 nanoclusters (Fig. 1B i-ii; Fig. 1C i-v). To investigate whether Syt1 undergoes co-clustering with cognate SV2 molecules, dual-colour imaging of Syt1^WT^-pH-atto647N-NB with mEos2-tagged wild-type SV2A (SV2A^WT^-mEos2) was performed. Syt1 and SV2 nanoclusters overlapped along hippocampal nerve terminals (Fig. 1B). To determine whether Syt1 interaction with SV2A regulates the nanoscale organisation of Syt1 at the PM, we expressed a pH-tagged Syt1 mutant that has perturbed SV2A binding (Syt1^K326A/K328A^-pH; Fig. 1A i, 1D i-v) (Borden et al., 2005). This Syt1 variant contains two lysine (K) to alanine (A) substitutions within the polybasic region of Syt1’s C2B domain. A loss of Syt1 subsynaptic clustering was observed upon expression of the K326A/K328A mutant (Fig. 1D iii-v), despite Syt1^WT^-pH and Syt1^K326A/K328A^-pH being expressed at comparable levels (Fig. 1E i).

**Figure 1.**
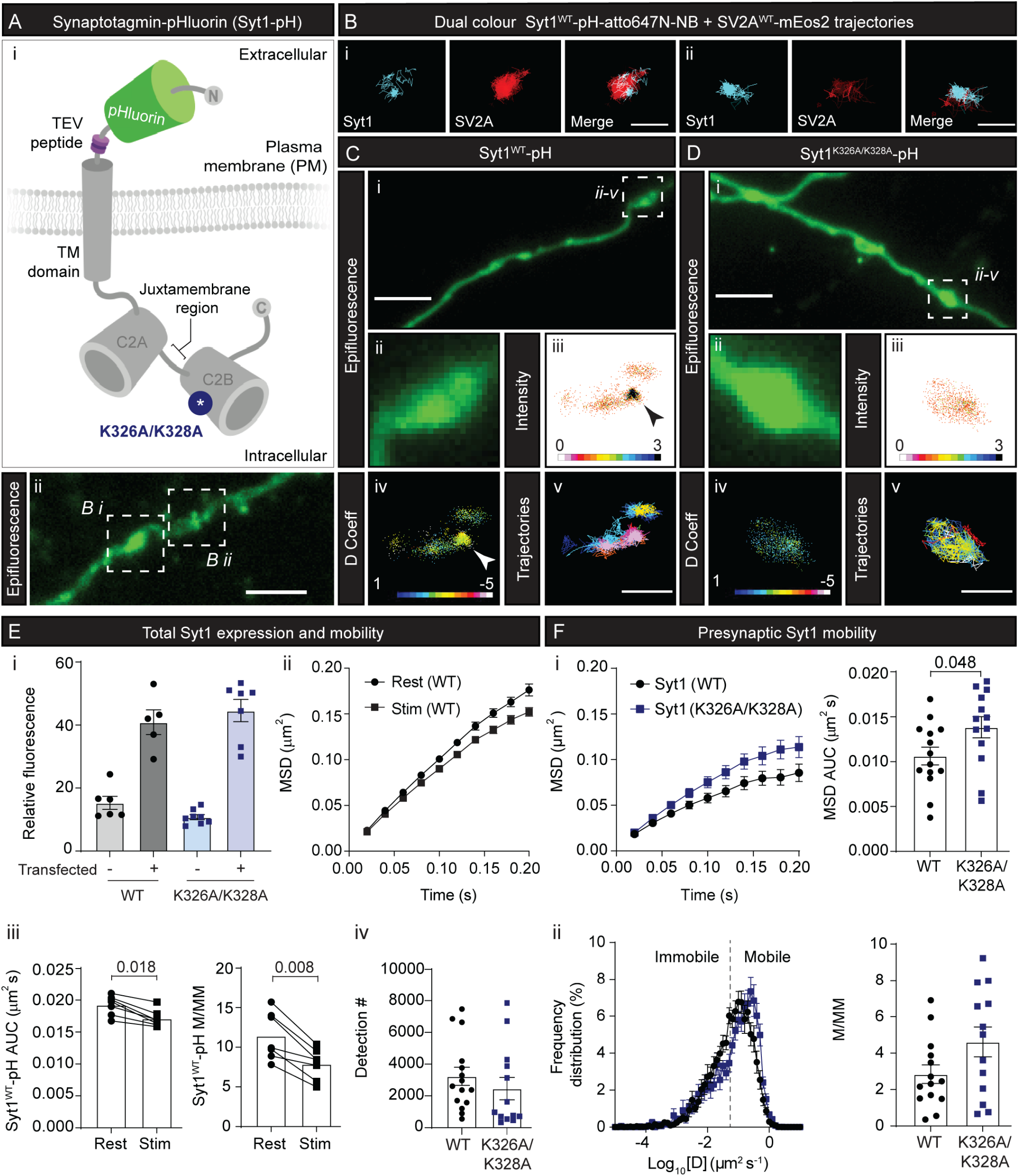
Inhibition of SV2A interaction (K326A/K328A) increases the mobility of Syt1 following stimulation. Universal Point Accumulation for Imaging in Nanoscale Topography (uPAINT) of Syt1-pH at the plasma membrane (PM) was performed in murine hippocampal neurons (DIV 18) treated with anti-GFP atto647N-nanobodies (NB). **(A)** (i) Schematic of Syt1-pH at the PM (post fusion). The K326A/K328A mutations (blue) are in the intracellular C2B domain of Syt1. The TEV peptide sequence (purple) and pHluorin tag (green) are shown. (ii) Syt1^WT^-pH epifluorescence in the axon. Scale bar = 5 µm. Regions of interest surrounded by white dotted outlines are shown magnified in B. **(B)** (i-ii) Clustered Syt1^WT^-pH-atto647N-NB trajectories (cyan) localizing within SV2A^WT^-mEos2 (red) hotspots. Scale bar = 1 µm. **(C)** Syt1^WT^-pH epifluorescence in (i) axons and (ii) nerve terminals. Super-resolved Syt1^WT^-pH-atto647N-NB in nerve terminals highlighted by (iii) intensity, (iv) diffusion coefficient (D Coeff) and (v) trajectory maps. For the D Coeff panel, the colour bar represents log_10_[D] (µm^2^s^-1^), with warmer colours indicating points of low mobility. Arrowheads designate Syt1^WT^-pH-atto647N-NB clusters. Syt1^K326A/K328A^-pH epifluorescence in (i) axon (ii) nerve terminals. Super-resolved Syt1^K326A/K328A^-pH-atto647N-NB highlighted by (iii) intensity, (iv) D Coeff and (v) trajectories. Axon scale bar = 4 µm (C i, D i), presynapse scale bar = 1 µm (C v, D v). **(E)** Total axonal Syt1 mobility. (i) Syt1 expression levels in neurons transfected with either Syt1^WT^-pH (untransfected control n=6; WT n=5) or Syt1^K326A/K328A^-pH (untransfected control n=8; K326A/K328A n=7), and subsequently fixed and stained using an anti-Syt1 antibody. (ii-iii) Single molecule mobility of Syt1^WT^-pH-atto647N-NB imaged in live neurons under resting and stimulated (high K^+^; 56 mM) conditions, with the corresponding (ii) mean square displacement (MSD; µm^2^ over 200 ms), (iii, left) the area under the curve (AUC) of MSD (µm^2^s), and (iii, right) the mobile-to-immobile ratio (M/MM) shown (n=7). (iv) Total surface detections of Syt1^WT^-pH and Syt1^K326A/K328A^-pH. **(F)** Presynaptic mobility of Syt1^WT^-pH-atto647N-NB (n=14) and Syt1^K326A/K328A^-pH-atto647N-NB (n=13), represented as (i) the MSD and corresponding AUC of the MSD, (ii) frequency distribution (%) using log_10_[D] (µm^2^s^-1^) values whereby D is diffusion coefficient, and the M/MM ratio. Statistical significance was determined using a Student’s *t* test. N values are based on the number of neurons.

To determine whether interaction with SV2A controlled the lateral confinement of Syt1 at the PM, the surface mobility of Syt1^WT^-pH and Syt1^K326A/K328A^-pH was examined. First, we assessed whether the mobility Syt1 was affected by synaptic activity, due to prominent clustering having been observed following stimulation. We tracked Syt1^WT^-pH-atto647N-NB single molecules before and after stimulation across the total hippocampal axon and plotted the mean square displacement (MSD) of Syt1^WT^-pH molecules over time (200 ms) (Fig. 1E ii). This was quantified by calculating the area under the curve (AUC) of the MSD, and by calculating the ratio of mobile-to-immobile molecules (M/MM) (Fig. 1E iii). Both metrics were significantly decreased following stimulation (p=0.018 and p=0.008 respectively). Further, we used an intraluminal Syt1 nanobody and found that the mobility of overexpressed and endogenous Syt1 were identical (data not shown). These results demonstrate that the lateral entrapment of Syt1 molecules at the PM is an activity-dependent process and not due to increased surface expression. Therefore, to assess the impact of preventing Syt1 from interacting with SV2A, we compared the surface mobility of Syt1^WT^-pH and Syt1^K326A/K328A^-pH, taking advantage of the unquenching of the pH-tag to identify active presynapses. To control for the impact of Syt1 surface expression on Syt1 mobility, the number of Syt1^WT^-pH and Syt1^K326A/K328A^-pH detections at the PM were quantified post-stimulation and found to be similar (Fig. 1E iv). As anticipated, Syt1^WT^-pH mobility at the presynapse (Fig. F i) was lower compared to that of the whole axon (Fig. 1E iii), suggesting that Syt1 is specifically confined at the presynaptic membrane. We next compared Syt1^WT^-pH and Syt1^K326A/K328A^-pH mobility by plotting their MSD and AUC of the MSD (Fig. 1F i), as well as their frequency distribution and M/MM ratio (Fig. 1F ii). Although the observed positive shift in the M/MM ratio for Syt1^K326A/K328A^-pH was not significant (p=0.07; Fig.1F ii), the MSD was significantly higher than that of Syt1^WT^-pH (p=0.048; Fig.1F i). These results suggest that the binding of Syt1 via its K326/K328 residues to SV2A controls its confinement at the PM in an activity-dependent manner.

### Knockdown of endogenous SV2A increases the surface mobility of Syt1 and impairs Syt1 nanocluster formation

To confirm that the observed increase in Syt1 surface mobility in response to the K326A/K328A mutation was specific to perturbed SV2A binding, we knocked down endogenous SV2A by transfecting neurons with an mCerulean (mCer) tagged SV2A-shRNA (SV2A-shRNA-mCer) (Fig. 2A) (Dong et al., 2006; Harper et al., 2020; Zhang et al., 1994; Zhang et al., 2015). As previously reported in these studies, a significant decrease (p<0.0001) in endogenous SV2A expression was observed in the presence of SV2A-shRNA-mCer (Fig. 2A-B). To determine if changes in surface mobility were specific to Syt1, another vesicular protein, VAMP2, was tracked (VAMP2-pH-atto647N-NB) in the presence of either SV2A-shRNA-mCer, or empty-vector mCer (Fig. 2C). We also tracked Syt1^WT^-pH-atto647N-NB in the presence of either empty-vector mCer, SV2A-shRNA-mCer (knock-down) or a bicistronic plasmid encoding both SV2A-shRNA and shRNA-resistant SV2A-mCer (knock-down rescue) (Fig 2D). Notably, the knockdown of endogenous SV2A increased the surface mobility of Syt1^WT^-pH-atto647N-NB at the presynapse, as evident by a significant increase in the AUC of the MSD (Fig. 2E), and a significant decrease in the percentage of immobile molecules (Fig. 2F). For rescued expression, Syt1^WT^-pH-atto647N-NB was tracked in cells where SV2A was re-expressed using a bicistronic plasmid that encoded SV2A-shRNA and shRNA-resistant SV2A-mCer (Fig 2D). This change in Syt1^WT^-pH-atto647N-NB mobility in the presence of the SV2A-shRNA was alleviated upon rescue of endogenous SV2A expression using the shRNA-resistant SV2A-mCer (Fig. 2E-F). In contrast to this, no change in mobility was observed for VAMP2-pH-atto647N-NB in the presence of the SV2A-shRNA knockdown (Fig. 2G-H), indicating that these effects were Syt1 specific. These results therefore suggest that SV2A plays a key role in sequestering Syt1 into nanoclusters at the PM. Although we cannot rule out that there are other proteins that also contribute to the trapping of Syt1 on the PM, the fact that VAMP2 mobility is unchanged following SV2A knockdown suggests that the entrapment effect of SV2A is specific to Syt1.

**Figure 2.**
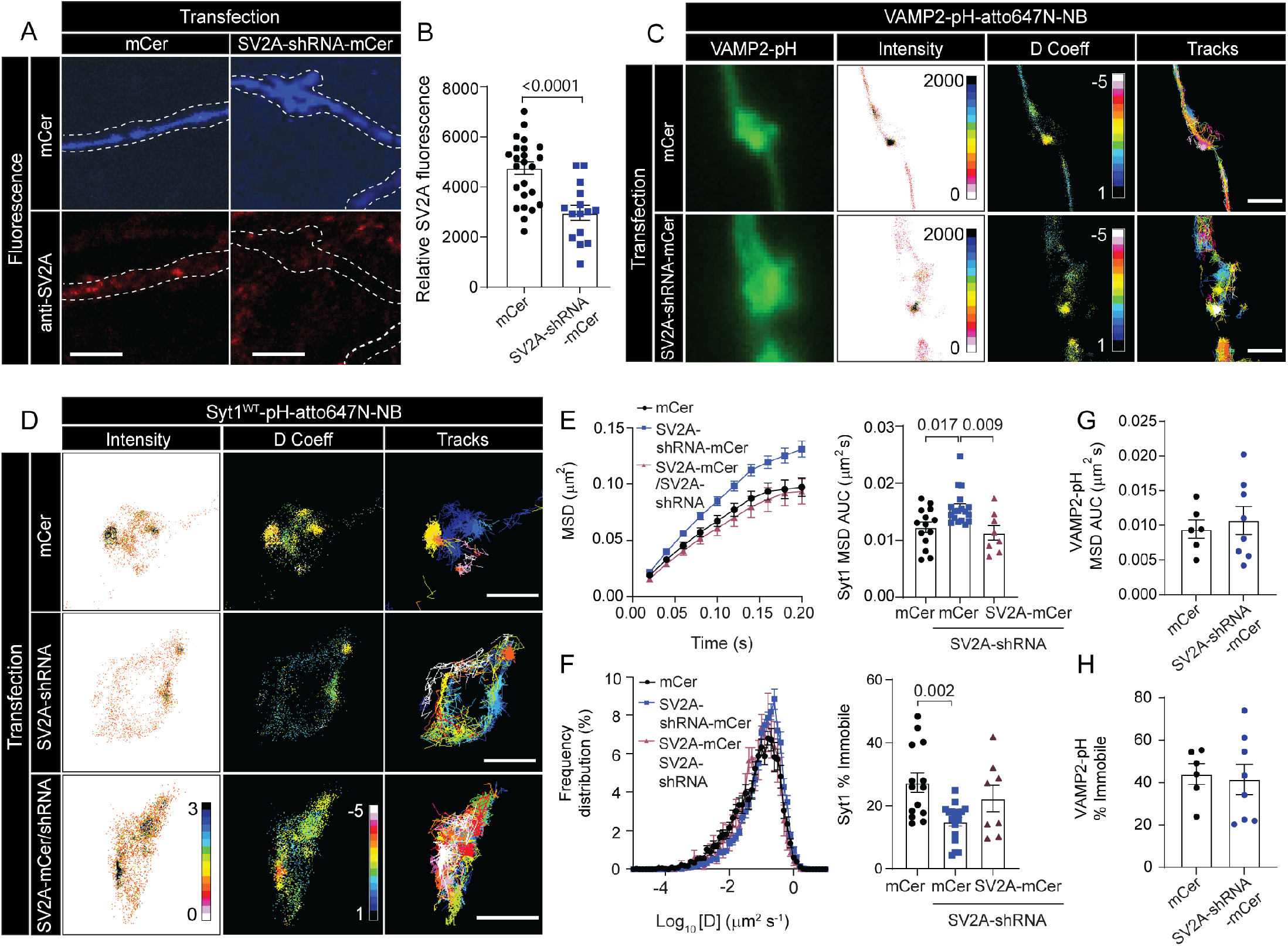
Depletion of SV2A increases Syt1 mobility which is rescued by SV2A-mCer re-expression. **(A)** Expression of either mCer or SV2A-shRNA-mCer with endogenous SV2 staining in hippocampal neurons. Axons of transfected neurons are highlighted by white dotted outlines. Scale bar = 10 µm. **(B)** Validation of SV2A knockdown, with SV2A expression measured using relative fluorescence in cells transfected with either SV2A-shRNA-mCer (knockdown; n=15) or mCer (knockdown control; n=24). **(C)** VAMP2-pH co-expressed with either mCer or SV2A-shRNA-mCer. VAMP2-pH epifluorescence is shown alongside the average intensity, diffusion coefficient and trajectory maps of VAMP2-pH-atto647N-NB. For the D Coeff panel, the colour bar represents log_10_[D] (µm^2^s^-1^), with warmer colours indicating points of low mobility. Scale bar = 1 µm. **(D)** Syt1^WT^-pH-atto647N-NB co-expressed in neurons with either mCer (knockdown control; n=14), SV2A-shRNA-mCer alone (knockdown; n=18) or SV2A-shRNA with an shRNA-resistant SV2A-mCer (knockdown rescue; n=8). Average intensity, diffusion coefficient and trajectory maps of Syt1^WT^-pH-atto647N-NB are shown. Scale bar = 1 µm. **(E)** MSD of Syt1^WT^-pH-atto647N-NB, and corresponding AUC of the MSD. **(F)** Frequency distribution of Log_10_ [D] values for Syt1^WT^-pH-atto647N-NB trajectories, and percentage of immobile Syt1^WT^-pH-atto647N-NB molecules. MSD of the AUC and **(H)** percentage of immobile VAMP2-pH-atto647N-NB molecules in the presence of either mCer (control; n=6) or SV2A-shRNA-mCer (knockdown; n=8). Statistical significance was determined using a one-way ANOVA and Tukey’s test for multiple comparisons, and Student’s *t*-test for single comparisons.

### Quantification of Syt1 nanoclustering at the plasma membrane

The increased mobility of Syt1 in the absence of SV2A expression or interaction suggests that SV2A plays a role in organising Syt1 into nanoclusters at the PM. Based on this, we first defined the dimensions of Syt1^WT^-pH nanoclusters using the newly-developed nanoscale spatiotemporal indexing clustering (NASTIC) analysis (Fig. 3A-D) (Wallis et al., 2021). Under physiological conditions, Syt1 nanoclusters were found to be distributed across the axonal branches of hippocampal neurons (Fig. 3A). To assess whether these Syt1 nanoclusters were associated with endocytic sites, we performed dual colour imaging of Syt1^WT^-pH-atto647N-NB using uPAINT in tandem with single-particle tracking photoactivated localization microscopy (sptPALM) of mEos4b-clathrin. This revealed that Syt1 nanoclusters were formed in regions devoid of clathrin, therefore suggesting that Syt1 nanoclustering at the PM occurs independently of clathrin-mediated endocytosis (Fig. 3B iii-v). Next, to determine whether the increased mobility of the Syt1 SV2A binding mutant was due an alteration in nanoclustering, we used cluster analysis to quantify the size, density, and apparent lifetime of both Syt1^WT^-pH and Syt1^K326A/K328A^-pH nanoclusters. By doing so, we identified a portion of Syt1^WT^-pH molecules that formed nanoclusters (4.68 ± 0.87 %). These nanoclusters had a mean area of 0.042 ± 0.0015 µm^2^ and were present for a short duration (6.18 ± 0.28 sec). Although the MSD (Fig. 3C i), detection frequency (Fig. 3C ii) and apparent lifetime (Fig. 3C iii) of Syt1^K326A/K328A^-pH within clusters was not significantly different compared that of Syt1^WT^-pH, there was a significant decrease in nanocluster density (p=0.048) (Fig. 3C iv), and an increase in both cluster area (p=0.012) (Fig. 3C v) and radius (p=0.008) (Fig. 3C vi). Similarly, knockdown of endogenous SV2A expression using SV2A-shRNA-mCer caused a significant reduction in the percentage of clustered Syt1^WT^-pH trajectories at the PM (p=0.002; Fig. 3D i). Furthermore, although there was only a marginal decrease in the number of trajectories per cluster of Syt1 following knockdown of SV2A (p=0.055; Fig. 3D ii), a significant decrease in both nanocluster area (p=0.006; Fig. 3D iii), and apparent cluster lifetime (p=0.035; Fig. 3D iv) was observed. Taken together, these results demonstrate that expression of (and interaction with) SV2A facilitates the entrapment of Syt1 within nanoclusters at the PM.

**Figure 3.**
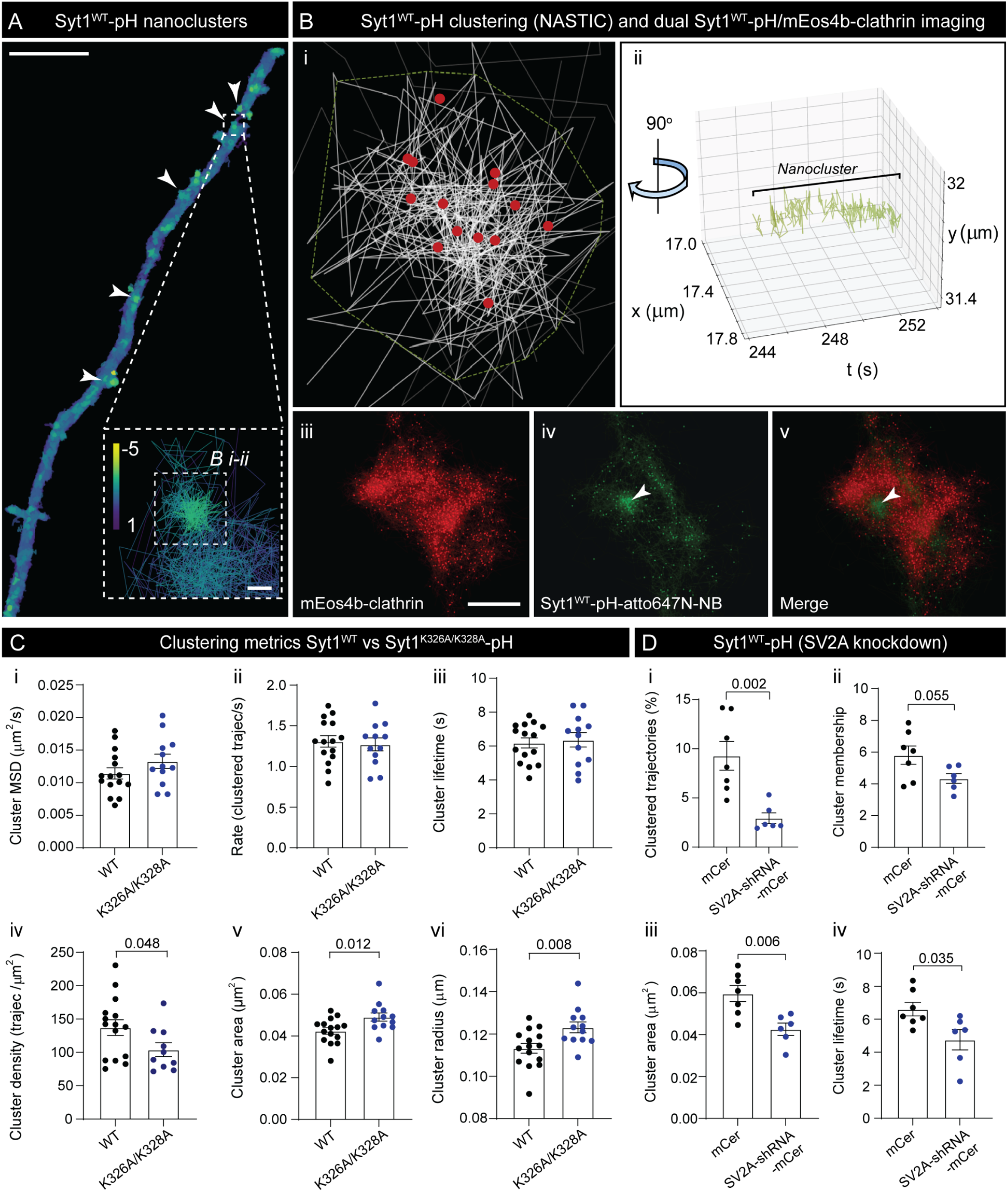
Syt1 nanoclusters are segregated from clathrin and are altered upon Syt1 K326A/K328A mutation and SV2 knockdown. **(A)** Syt1-pH-atto647N-NB nanoclusters on the PM, indicated by white arrowheads. Insert shows example nanocluster trajectories (analysed in B i-ii) with variable D Coeff values (Log_10_[D] from -5 to 1 (um^2^s^-1^); warmer colours indicate points of low mobility. Scale bar = 5 µm (axon), and 0.1 µm (zoom). **(B)** Spatiotemporal alignment (xyt) of Syt1^WT^-pH nanoclusters in the presynapse, and segregation from mEos4b-clathrin clusters. (i) Enhanced view of a single Syt1^WT^-pH nanocluster shown in A, with tracks (white), boundary (green dotted outline) and centroids (red). (ii) Lifetime of the nanocluster shown in B i (tracks rotated ∼90° across xyt) obtained using Nanoscale Spatiotemporal Indexing Clustering (NASTIC). Trajectory centroids of (iii) mEos4b-clathrin and (iv) Syt1^WT^-pH-atto647N-NB were (v) merged to show exclusion of Syt1 nanoclusters (green) from clathrin (red). Syt1 nanocluster indicated by arrowheads. Scale bar = 0.5 µm. **(C)** Nanocluster metrics of Syt1^WT^-pH and Syt1^K326A/K328A^-pH (WT n=15 neurons and K326A/K328A n=12 neurons), quantified using (i) AUC of the MSD, (ii) rate of clustering (trajectories/s), (iii) apparent lifetime (s), (iv) density (trajectories/µm^2^), (v) area (µm^2^) and (vi) radius (µm). **(D)** Nanocluster metrics of Syt1^WT^-pH following SV2A knockdown (mCer n=7 neurons; SV2A-shRNA-mCer n=6 neurons), quantified by (i) the percentage of clustered trajectories, (ii) cluster membership (trajectories/cluster), (iii) area and (iv) apparent lifetime. Statistical significance was determined using a Student’s *t-*test.

### SV2A mobility depends on binding to Syt1 but not to endocytic machinery

Our results demonstrate that the organisation of Syt1 into nanoclusters depends on its interaction with SV2A, and that these Syt1 nanoclusters do not co-localise with clathrin clusters at the PM. Based on this hypothesis, we next investigated whether SV2A nanoclusters follow similar dynamics. To this end, we imaged a Syt1-binding SV2A mutant (SV2A^T84A^), and an SV2A mutant that lacks binding to clathrin adaptor AP2 (SV2A^Y46A^) (Yao et al., 2010), and examined their nanoscale mobility (Fig. 4A). To validate that binding of Syt1 to SV2A^T84A^ was perturbed, we co-immunoprecipitated SV2A from HEK-293T cells co-expressing Syt1-HA with either SV2A^WT^-mCer, SV2A^T84A^-mCer or SV2A^Y46A^-mCer, by use of an anti-GFP antibody which also recognises ECFP derivative mCer (Fig. 4B). The level of Syt1-HA pulldown was significantly reduced for SV2A^T84A^-mCer (Fig. 4B-C). In contrast to this, the opposite effect was observed for SV2A^Y46A^-mCer AP2-binding mutant, with increased Syt1-HA binding detected (Fig. 4B-C). This finding suggests that AP2 and Syt1 compete for binding to SV2A, likely due to the proximity of the Syt1 (T84) and AP2 (Y46) interaction sites (Fig. 4A). Next, we performed uPAINT imaging of hippocampal neurons expressing either SV2A^WT^-pH, SV2A^T84A^-pH, or SV2A^Y46A^-pH. SV2A^WT^-pH and SV2A^Y46A^-pH formed nanoclusters within presynaptic boutons, which were less prominent for SV2A^T84A^-pH (Fig. 4D). Cluster analysis revealed a decrease in the apparent lifetime of SV2A^T84A^-pH nanoclusters compared to that of SV2A^WT^-pH (Fig. 4D i). However, nanocluster area and membership were unchanged (Fig. 4D ii-iii). Further, SV2A^T84A^-pH mobility was significantly increased in comparison to SV2A^WT^-pH (Fig. 4E i-ii), demonstrating that the SV2A-Syt1 interaction is essential for the trapping of both molecules at the PM. Loss of AP2 binding (SV2A^Y46A^-pH) however, had no effect on the surface mobility of SV2A (Fig. 4E i-ii), indicating that nanoclustering of SV2A is not regulated by the endocytic machinery. This was further confirmed by treating cells with Dyngo4A (30 μM for 30 min) – an inhibitor of the GTPase activity of dynamin (McCluskey et al., 2013), which failed to elicit a change in the MSD of SV2A^WT^-pH (Fig. 4E iii-iv). Overall, our results suggest that the nanoclustering of Syt1 and SV2A on the PM is controlled by their mutual direct interaction and is independent from their binding to endocytic machinery.

**Figure 4.**
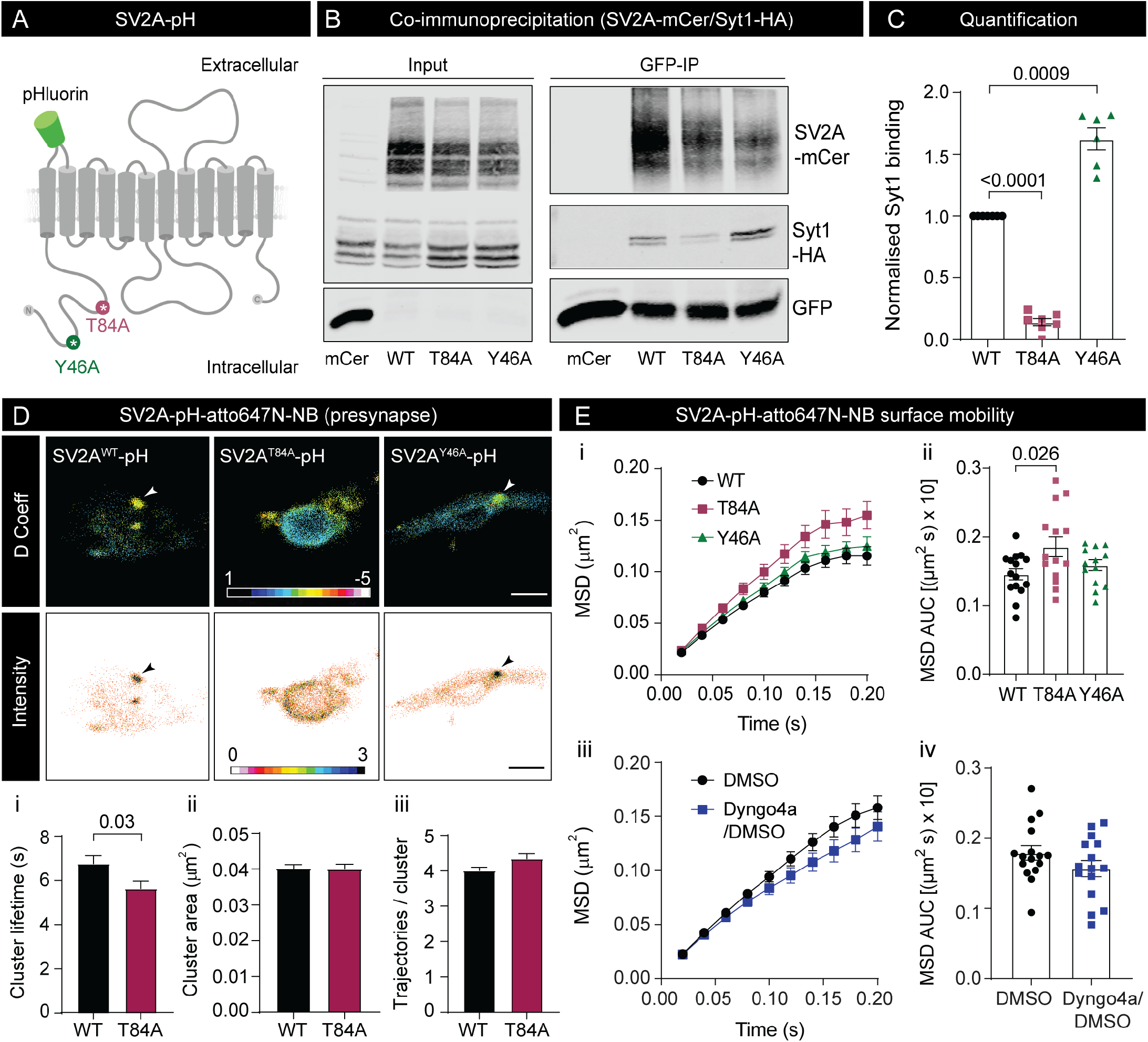
SV2A nanoclustering is controlled by Syt1 interaction (T84A), and unaffected by AP2 interaction (Y46A) and dynamin inhibition. **(A)** Schematic of SV2A-pH at the PM (post fusion). The pH tag (green) is located between the first and second transmembrane domains. Note the proximity of the N-terminal T84A (red) and Y46A epitopes (green). **(B)** Co-immunoprecipitation of Syt1-HA with either mCer (control), SV2A^WT^-mCer, SV2A^T84A^-mCer or SV2A^Y46A^-mCer from protein samples derived from HEK-293T cells, using anti-GFP antibody conjugated TRAP beads. Representative blots using total protein lysates (input), and GFP immunoprecipitation of SV2A-mCer (GFP-IP), are shown. **(C)** Level of Syt1-HA binding by either SV2A^T84A^-mCer or SV2A^Y46A^-mCer as normalised to that of SV2A^WT^-Cer (WT and T84A: n=7, Y46A: n=6 samples per group). **(D)** Super-resolved SV2A-pH-atto647N-NB (WT, T84A or Y46A) within the presynapse of hippocampal nerve terminals. For D Coeff panels, regions highlighted in warm colours represent points of low mobility. Arrowheads indicate points of nanoclustering. Scale bar = 1 µm. (i) Nanocluster apparent lifetime, (ii) area and (iii) membership of SV2A^WT^-pH-atto647N-NB (n=221 clusters) and SV2A^T84A^-pH-atto647N-NB (n=177 clusters). **(E)** Surface mobility of SV2A-pH-atto647N-NB (SV2A^WT^ n=15; SV2A^T84A^ n=14; SV2A^Y46A^ n=13) at the presynapse, as assessed by (i) MSD and (ii) AUC of the MSD (µm^2^ s × 10). (iii-iv) SV2A^WT^-pH-atto647N-NB mobility following treatment with either DMSO (control; n=16), or Dyngo4A in DMSO (30 µM for 30 min; n=15), as assessed by (iii) MSD and (iv) AUC of the MSD (µm^2^ s × 10). Statistical significance was determined by a one-way ANOVA with a Tukey’s test for multiple comparisons and a Students’ *t*-test for single comparisons.

### Activity-dependent retrieval of SV2A is controlled by AP2 but not Syt1

To determine whether the clustering of SV2A is associated with alterations in SV endocytosis, we next examined the kinetics of SV2A-pH retrieval. For this, we expressed either SV2A^WT^-pH, SV2A^T84A^-pH or SV2A^Y46A^-pH in hippocampal neurons and measured the fluorescence decay of the pH-tag following electrical field stimulation (300 action potentials (APs) at 10 Hz), prior to treating neurons with ammonium chloride (NH_4_Cl) (Fig. 5A). The loss of pH fluorescence is reflective of SV2A-pH retrieval kinetics during SV endocytosis, which is rate limiting in comparison to subsequent SV acidification (Atluri and Ryan, 2006; Granseth et al., 2006). Rapid de-acidification of SVs following NH_4_Cl treatment results in the total population of SV2A-pH becoming unquenched. These experiments revealed that retrieval of SV2A^Y46A^-pH was compromised, suggesting that interactions with AP2 are required for its efficient recovery during endocytosis (Fig. 5B i-ii). In comparison to this, impairing the interaction of SV2A with Syt1 (SV2A^T84A^-pH) did not have an impact on SV2A retrieval (Fig. 5B i-ii). As there were no differences in the proportion of total SV2A-pH following NH_4_Cl treatment between SV2A^WT^-pH, SV2A^T84A^-pH and SV2A^Y46A^-pH, the observed delay in SV2A^Y46A^-pH retrieval was not due to alterations in SV exocytosis (Fig. 5B iii). These findings demonstrate that AP2-mediated endocytosis of SV2A-pH occurs independently of SV2A-Syt1 interaction and nanoclustering.

**Figure 5.**
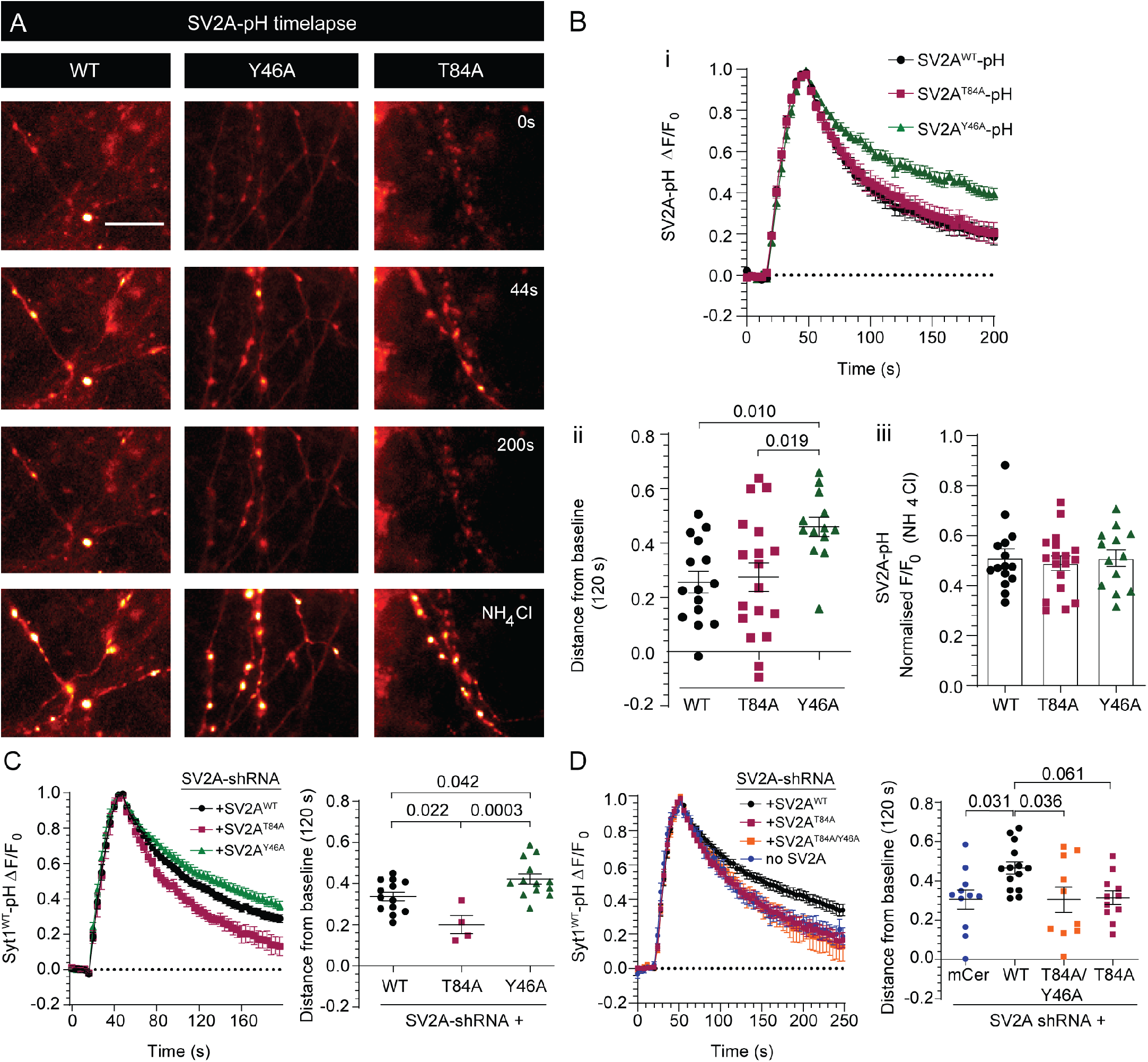
Activity-dependent endocytosis of Syt1 is controlled by SV2A. **(A)** Timelapse of SV2A^WT^-pH, SV2A^Y46A^-pH and SV2A^T84A^-pH surface fluorescence following stimulation (10 Hz; 300 action potentials (APs); 0s, 44s and 200s), prior to treatment with ammonium chloride (NH_4_Cl). Scale bar = 20 µm. **(B)** (i) SV2A-pH retrieval time course as measured by changes in SV2A-pH fluorescence (F/F_0_) (SV2A^WT^ n=15; SV2A^T84A^ n=18; SV2A^Y46A^ n=13). (ii) Surface stranded SV2A-pH remaining to be retrieved (distance from baseline) at 120 s post-stimulation. (iii) Normalised F/F_0_ (NH_4_Cl perfusion). **(C)** Syt1^WT^-pH retrieval time course in SV2A knockdown neurons (SV2A-shRNA) in the presence of either shRNA-resistant SV2A^WT^-mCer (n=13), shRNA-resistant SV2A^T84A^-mCer (n=4) or shRNA-resistant SV2A^Y46A^-mCer (n=13) expression after stimulation (10 Hz, 300 AP), quantified by the distance from baseline at 120 s. **(D)** Time course of Syt1^WT^-pH fluorescence intensity in the presence of SV2A-shRNA alone (no SV2A; n=11), or SV2A-shRNA in the presence of either shRNA-resistant SV2A^WT^-mCer (n=14), shRNA-resistant SV2A^T84A^-mCer (n=11) or shRNA-resistant SV2A^T84A/Y46A^-mCer (n=10) following stimulation (10 Hz, 300 AP), quantified using distance from baseline at 120 s. Statistical significance was determined with Kruskal-Wallis test with Dunn’s test for multiple comparisons.

### SV2A controls the activity-dependent endocytosis of Syt1

Knockdown of SV2A and disruption of SV2A binding accelerates the internalisation of Syt1-pH during SV endocytosis (Harper et al., 2020; Kaempf et al., 2015; Zhang et al., 2015). In order to determine the effect of SV2A on Syt1-pH retrieval, we knocked down endogenous SV2A in neurons using bicistronic plasmids that co-express SV2A-shRNA with either SV2A^WT^-mCer (knockdown rescue), SV2A^T84A^-mCer (Syt1 binding mutant) or SV2A^Y46A^-mCer (AP2 binding mutant), which we co-transfected with Syt1^WT^-pH. By doing so, we confirmed that disruption of the SV2A-Syt1 interaction (SV2A^T84A^-mCer) accelerated the retrieval kinetics of Syt1^WT^-pH from the PM (Fig. 5C). Conversely, blocking the interaction between SV2A and AP2 (SV2A^Y46A^-mCer) slowed the activity-dependent retrieval of Syt1^WT^-pH (Fig. 5C). To assess whether the accelerated retrieval of Syt1^WT^-pH that was observed in response to SV2A^T84A^-mCer expression was due to lack of competition with AP2, we also performed the SV2A knockdown experiment in the presence of a SV2A double mutant that has both perturbed Syt1 and AP2 binding (SV2A^T84A/Y46A^-mCer) (Fig. 5D). As an additional control, we also performed SV2A knockdown in the absence of the SV2A rescue construct (SV2A-shRNA alone: no SV2A) and assessed the effect of Syt1^WT^-pH retrieval kinetics. The rate of Syt1^WT^-pH internalisation was not rescued by the additive loss of AP2 binding (Fig. 5D), indicating that SV2A interaction is the dominant mechanism through which Syt1 endocytosis is regulated.

### SV2A knockdown increases dynamin1 recruitment at the plasma membrane

The accelerated endocytosis of Syt1 in the absence of SV2A suggests that compensatory recruitment of endocytic machinery is taking place. For this reason, we examined the effect of perturbing SV2A expression on dynamin1 recruitment. Due to the reduced surface area and thin architecture of neurons, which would impede interpretation of these results, these experiments were instead conducted in pheochromocytoma (PC12) neurosecretory cells which have a larger surface area and have previously been used for SV2A-shRNA knockdown (Dong et al., 2006). Using TIRF microscopy, we conducted time-lapse imaging of GFP-tagged dynamin1 (Dyn1-GFP) (Xue et al., 2011). We analysed the effect of SV2A knockdown on the activity-dependent recruitment of dynamin1 to the PM, by transfecting PC12 cells with either SV2A-shRNA-mCer (knockdown) or empty vector mCer (knockdown control; Fig. 6A-C). Stimulation of PC12 cells with BaCl (2 mM) resulted in a substantial increase in dynamin1 recruitment to the PM, as indicated by a significant increase in Dyn1-GFP fluorescence intensity (FI; Fig. 6B-C). Importantly, this effect was potentiated by knocking down SV2A (Fig. 6B-C). Subsequently, we assessed the effect of SV2A knockdown on the clustering and mobility of dynamin1 on the PM at the presynapse, by performing sptPALM on hippocampal neurons co-expressing Dyn1-mEos2 with either mCer or SV2A-shRNA-mCer (Fig 6D-F). Knockdown of endogenous SV2A led to an increase in synaptic clustering (Fig. 6D) as evidenced by a large decrease in the AUC of the MSD of Dyn1-mEos2 (Fig. 6E), and a shift in the M/MM ratio of Dyn1-mEos2 towards decreased mobility (Fig. 6F). These findings suggest that increased dynamin1 recruitment occurs in response to de-clustering of stranded Syt1 at the PM. To determine whether these effects also occurred for Syt1 in the absence of dynamin1 interaction, the mobility of a phospho-inhibitory mutant of Syt1 tagged with pH (Syt1^T112A^-pH) was examined. This mutation is present within the juxtamembrane region of Syt1 (between the C2 domains; Fig. 6G i) and has been reported to decrease interaction with dynamin1 (De Jong et al., 2016; McAdam et al., 2015). No change in the surface mobility of Syt1 in either the axons or nerve terminals of hippocampal neurons was observed for this dynamin1-binding Syt1 mutant (Fig 6H-I). Further, pharmacological inhibition of dynamin’s enzymatic activity using Dyngo4A treatment (30 µM for 30 min) also failed to elicit a change in Syt1 mobility (Fig. 6I). Coupled with the previous observation that SV2A mobility was unaffected by AP2 interaction and dynamin inhibition (Fig. 4), these results suggest that the endocytic machinery has a limited role in regulating the nanoscale organization of Syt1 and SV2A at the PM. Rather, Syt1-SV2 nanoclustering acts upstream of endocytosis by controlling the recruitment of the endocytic machinery.

**Figure 6.**
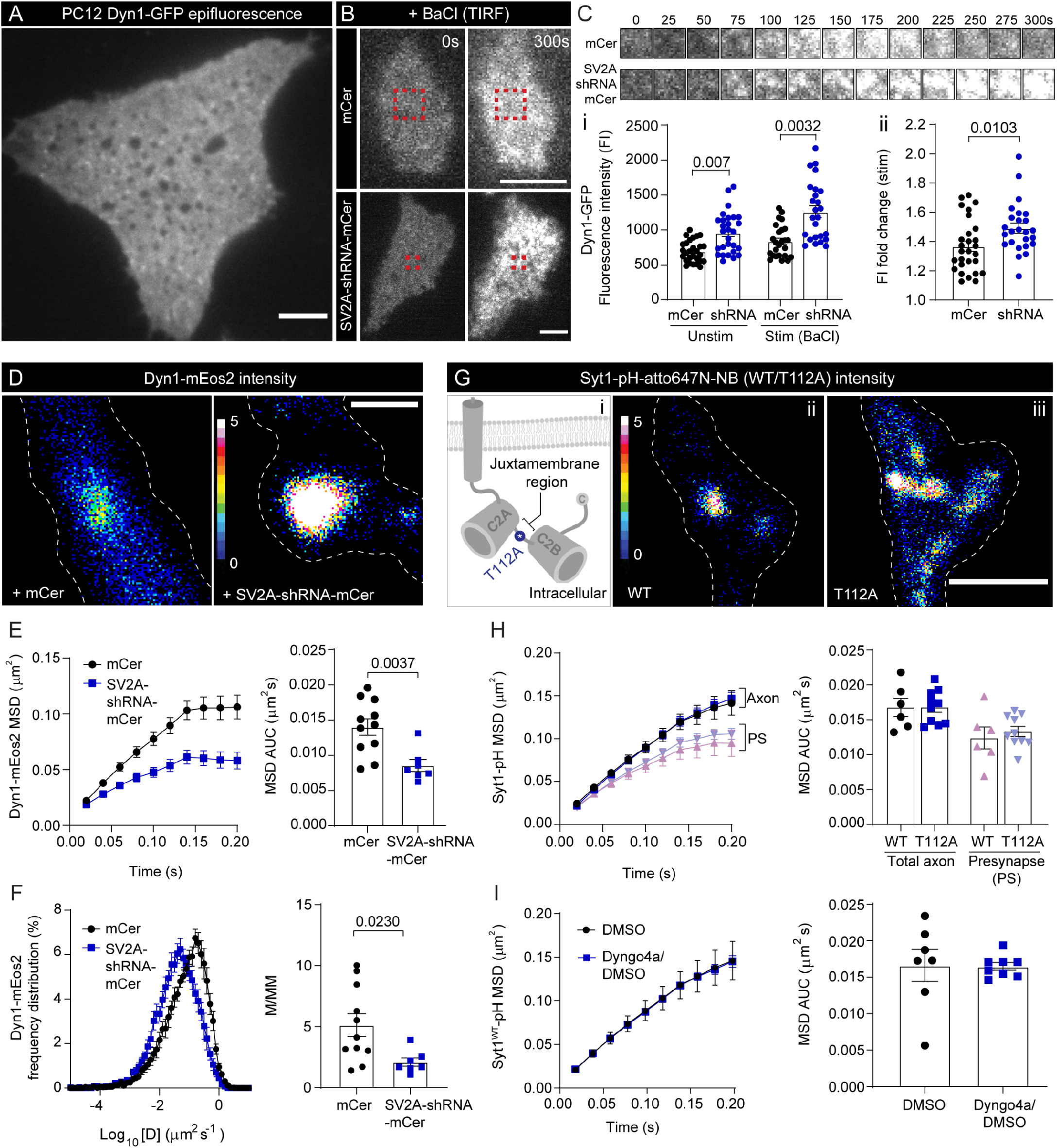
Dynamin1 activity-dependent recruitment to the PM is potentiated by SV2A knockdown and dynamin inhibition does not impact Syt1 mobility. **(A)** Dyn1-GFP epifluorescence in live PC12 cells. **(B)** Total Internal Reflection Fluorescence (TIRF) of Dyn1-GFP in the presence of either mCer (knockdown control) or SV2A-shRNA-mCer (knockdown) pre- and post-stimulation with BaCl (2 mM). Scale bar = 4 µm for A and B. **(C)** Timelapse of Dyn1-GFP in the presence of either mCer or SV2A-shRNA-mCer following BaCl stimulation (0-300 s). (i) Dyn1-GFP fluorescence intensity (FI) is quantified in the presence of either mCer or SV2A-shRNA-mCer both at rest and following BaCl stimulation. (ii) Fold change in Dyn1-GFP FI following stimulation. **(D)** Hippocampal neurons (DIV 18), co-transfected with Dyn1-mEos2 and either mCer or SV2A-shRNA-mCer, were imagined by sptPALM in TIRF. Average intensity showing clusters of Dyn1-mEos2 on the PM. Scale bar = 0.5 µm. Dyn1-mEos2 mobility in the indicated conditions quantified using **(E)** MSD and AUC of the MSD, and **(F)** Frequency distribution and M/MM ratio. **(G)** Hippocampal neurons were transfected with either Syt1^WT^-pH or Syt1^T112A^-pH (a dynamin binding deficient mutation) and imaged with uPAINT using atto647N-NB following high K^+^ stimulation. (i) Schematic highlighting the T112A mutation (blue) within the juxtamembrane domain of Syt1. Average intensity of (ii) Syt1^WT^-pH-atto647N-NB and (iii) Syt1^T112A^-pH-atto647N-NB. White dotted outline of nerve terminal area. Scale bar =1 µm. **(H)** Quantification of Syt1^WT^-pH-atto647N-NB mobility in both the axon and presynapse (PS) as shown by the MSD and the AUC of the MSD. No significant difference in mobility was observed between Syt^WT^-pH and Syt1^T112A^-pH in either the axon or presynapse (PS). **(I)** Syt1^WT^-pH-atto647N-NB mobility in the presence of either DMSO (control) or the dynamin inhibitor Dyngo4a (30 mM, 30 min), shown as MSD and AUC of the MSD. Statistical significance was determined using Student’s *t* test.

### Loss of SV2A interaction alters the intracellular sorting of Syt1 at the recycling pool of SVs

The accelerated retrieval of Syt1 that was observed in the presence of the Syt1-binding SV2A mutant (SV2A^T84A^) raises the question of whether SV2A controls the endocytic targeting of Syt1 to recycling SVs. If this is the case, interfering with SV2A may also causes intracellular missorting of Syt1. To address this question, we performed the sub-diffractional Tracking of Internalised Molecules (sdTIM) technique (Joensuu et al., 2017; Joensuu et al., 2016) which enables the imaging of internalised Syt1. Syt1-pH-positive neurons were pulsed (56 mM K^+^ with anti-GFP atto647N-NB for 5 min) before being washed and chased under resting conditions (5.6 mM K^+^ for 5 min). To selectively image the recycling pool of SVs containing Syt1 with greater accuracy, we cleaved the Tobacco-Etch Virus (TEV) sequence present between Syt1 and the pH tag, using an active TEV protease (60 U for 15 min), thereby removing the pH-atto647N-NB tag from Syt1-pH-atto647N-NB that remained stranded at the surface (Fig. 7A i-iii) (Gimber et al., 2015; Hua et al., 2011; Nair et al., 2013; Wienisch and Klingauf, 2006). As expected, we observed a decrease in Syt1^WT^-pH and Syt1^K326A/K328A^-pH fluorescence in the presence of the active TEV protease compared to the inactivated boiled TEV control (95 °C for 10 min; Fig. 7A iv) and were able to track distinct recycling SVs containing Syt1-pH-atto647N-NB across both the total axon and in nerve terminals following active TEV digestion (Fig. 7B). As a control, we assessed Syt1^WT^-pH and Syt1^K326A/K328A^-pH mobility in the presence of either inactive (Fig. 7C) or active (Fig. 7D) TEV protease in both the total axon (Fig. 7C i-ii and 7D i-ii) and at the presynapse (Fig. 7C iii-iv and 7D iii-iv). In the absence of surface digestion, Syt1^WT^-pH and Syt1^K326A/K328A^-pH had a comparable mobility across both the entire axon and within nerve terminals (Fig. 7C). This mobility was reduced upon surface digestion with active TEV (Fig. 7C-D), confirming that Syt1-pH-atto647N-NB was specifically imaged within the recycling pool of SVs. Although Syt1^WT^-pH and Syt1^K326A/K328A^-pH had similar mobilities across the whole axon following surface digestion (Fig. 7D i-ii), surprisingly, Syt1^K326A/K328A^-pH was significantly more mobile than Syt1^WT^-pH at the presynapse (Fig. 7D iii-iv). This suggests that loss of SV2A binding causes intracellular missorting of Syt1 specifically in nerve terminals. It is unlikely that the observed difference in mobility between Syt1^WT^-pH and Syt1^K326A/K328A^-pH is due to stranding of Syt1-pH at the PM, as no significant differences were observed in the absence of digestion of the surface fraction of Syt1-pH. Therefore, loss of SV2A binding may cause Syt1 to enter an alternative endocytic compartment, leading to differences in intracellular sorting.

**Figure 7.**
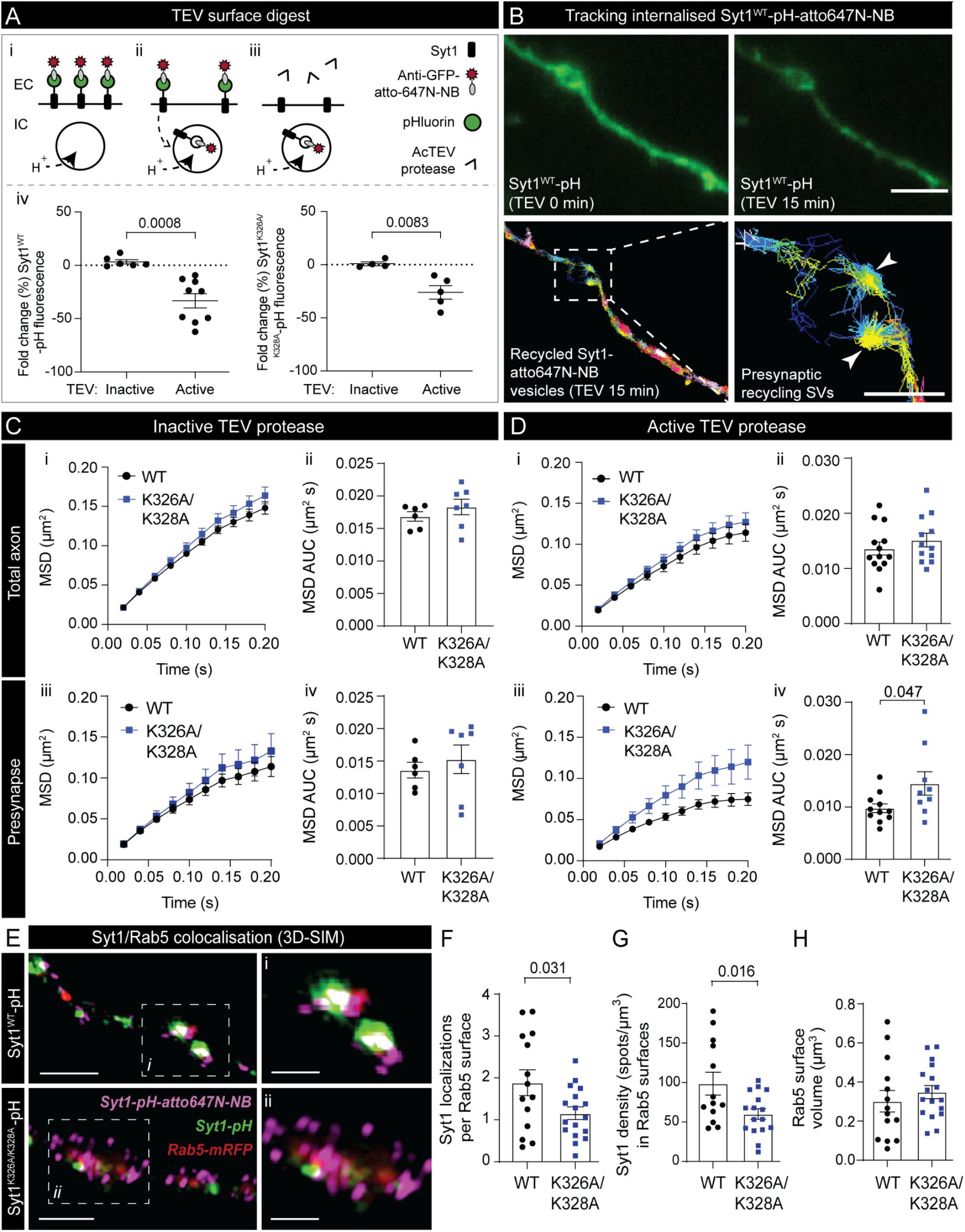
Syt1 intracellular sorting is altered in the absence of SV2A interaction. **(A)** Schematic depicting the removal of the pH-atto647N-NB tag from the surface fraction of Syt1-pH-atto647N-NB via active TEV protease digestion (60 U; 15 min), which enables the specific imaging of the internalised fraction of Syt1-pH-atto647N-NB. (i) Atto647N-NB binds to Syt1-pH present on the PM during stimulation (high K^+^; 5 min). (ii) Syt1-pH-atto647N-NB is internalised following a chase step (low K^+^; 5 min) into acidic vesicles which quenches the fluorescence of the pH tag. (iii) Active TEV protease cleaves pH-atto647N-NB from the remaining Syt1 surface fraction. (iv) Fold change (%) in Syt1-pH fluorescence in the presence of the inactive (boiled) and active TEV protease for Syt1^WT^-pH (n=6 inactive, n=9 active) and for Syt1^K326A/K328A^-pH (n=4 inactive, n=5 active). **(B)** Loss of Syt1^WT^-pH surface fluorescence induced by active TEV protease which allows selective tracking of internalised Syt1^WT^-pH-atto647N-NB within the recycling pool of SVs. Arrowheads indicate the trajectories of individual Syt1-positive recycling vesicles in a nerve terminal. Scale bar = 4 µm (axon) and 2 µm (presynapse). **(C-D)** Mobility of Syt1^WT^-pH-atto647N-NB and Syt1^K326A/K328A^-pH-atto647N-NB in the presence of **(C)** inactive and **(D)** active TEV, with the MSD and AUC of the MSD shown for (i-ii) the total axon and (iii-iv) the presynapse. **(E)** Hippocampal neurons were co-transfected with Rab5-mRFP and either Syt1^WT^-pH (n=14) or Syt1^K326A/K328A^-pH (n=17), fixed and imaged by 3D-structural illumination microscopy (3D-SIM). Scale bar = 2 µm (axon), 0.1 µm (zoom). Quantification showing **(F)** the number of Syt1-pH-atto647N-NB localizations per Rab5-positive vesicle surface, **(G)** Syt1-pH-atto647N-NB density within Rab5-mRFP surfaces (#spots/µm^3^), and the volume of Rab5-mRFP colocalised with Syt1-pH-atto647N-NB (µm^3^). Statistical significance was determined using Student’s *t* test.

### Loss of interaction with SV2A alters Syt1 trafficking, by sequestering it from Rab5-endosomes

Syt1 has previously been shown to be localized in early and recycling endosomes following internalisation 20 minutes post-fusion (Diril et al., 2006). It is becoming increasingly apparent that at physiological temperatures, both clathrin-dependent and clathrin-independent cargo sorting occurs at the level of internalised endosomes (Ivanova et al., 2021; Kononenko et al., 2013; Watanabe et al., 2014). Therefore, differences between Syt1^WT^-pH and Syt1^K326A/K328A^-pH mobility upon internalisation may stem from mutant or wild-type Syt1 being either sorted back to the recycling SV or to the endo-lysosomal system. To address this, we examined the co-localisation of internalised Syt1^WT^ or Syt1^K326A/K328A^ with Rab5 – a GTPase associated with early endosomes (Bucci et al., 1992) and bulk endosomes (Kokotos et al., 2018). Synaptotagmin has previously been shown to cluster within Rab5-positive early endosomes following internalisation (Hoopmann et al., 2010). Neurons were co-transfected with Rab5-mRFP (Vonderheit and Helenius, 2005) and either Syt1^WT^-pH or Syt1^K326A/K328A^-pH, and imaged using 3D structured illumination microscopy (3D-SIM) following activity-dependent internalisation of Syt1-pH-atto647N-NB (Fig. 7E). We characterised populations of Rab5-mRFP surface clusters that encased internalised Syt1-pH-atto647N-NB molecules. 3D-SIM revealed that a higher proportion of Syt1^WT^-pH-atto647N-NB localisations were identified in 3D-generated Rab5 surfaces as compared to Syt1^K326A/K328A^-pH-atto647N-NB (p=0.031) (Fig. 7F) Further, the volumetric density of Syt1^WT^-pH-atto647N-NB within these Rab5-mRFP surfaces was also significantly higher compared to the K326A/K328A mutant (p=0.016) (Fig. 7G). However, no significant difference in the endosomal surface volume was observed between Syt1^WT^-pH and mutant Syt1^K326A/K328A^-pH (Fig. 7H). Based on these findings, we concluded that inhibiting Syt1’s nanoclustering with SV2A decreased Syt1 intracellular trafficking towards Rab5-positive compartments (Fig. S2).

## Discussion

Interactions between vesicular proteins on the PM have recently been proposed to help maintain the organisation, stoichiometry, and composition of SVs by enhancing the fidelity of endocytic events (Gordon and Cousin, 2013; Harper *et al*., 2021; Harper *et al*., 2017). In this study, we provide evidence that the nanoclustering of SV2A-Syt1 at the PM controls the targeting of stranded vesicular proteins into recycling SVs. We demonstrate that SV2A controls the nanoclustering of Syt1 at the PM through interactions between its cytoplasmic N-terminus and the K326/K328 epitopes within the polybasic region of the C2B domain of Syt1. Importantly, this mechanism also works in reverse – SV2A is also sequestered into nanoclusters through its interaction with Syt1. On the other hand, genetic manipulation of the interaction with the endocytic machinery (AP2 and dynamin), as well as pharmacological inhibition of endocytosis (Dyngo4a), have no impact on either SV2A surface mobility or nanocluster formation. Syt1-SV2A co-clustering acts upstream of endocytosis and controls the kinetics of Syt1 endocytosis. This was made evident by the increase in recruitment to the PM, and immobilisation of dynamin1 in response to SV2A-Syt1 cluster inhibition. Therefore, this negative regulation of the endocytic machinery by SV2A-Syt1 nanocluster formation ultimately controls the rate and selective targeting of these surface stranded SV proteins into recycling SVs.

### Mechanisms regulating SV2A-Syt1 nanoclustering

The finding that SV2A controls Syt1 surface nanoclustering through interaction with the K326/K328 residues of Syt1 fits with previous observations that mutating these residues to alanines decreases Syt1 oligomerisation (Chapman et al., 1998). Here, we found that the reverse is also true – whereby SV2A nanoclustering is regulated via its interaction with these Syt1 residues. This is similar to what has been shown for other presynaptic molecules, such as syntaxin1A (Bademosi et al., 2016; Lang et al., 2001; Sieber et al., 2006) and Munc18 (Kasula et al., 2016), which form nanoclusters dependent on molecular interactions and play key roles in exocytosis. Importantly, SV2A-Syt1 nanoclustering is not regulated by the endocytic machinery (AP2 or dynamin1), unlike VAMP2 which does form endocytic machinery-mediated nanoclusters (Gimber et al., 2015). Vesicular proteins are therefore pre-assembled into nanoclusters on the PM following exocytosis, either by direct binary interaction or via interaction with the endocytic machinery. This process mediates the fidelity of their re-uptake into recycling SVs. In the case of SV2A, this hypothesis is supported by the observation that Syt1 and AP2 compete for interaction with SV2A, and that the endocytic kinetics of SV2A is unaffected by Syt1 interaction (SV2A^T84A^) and is instead dependent on AP2 binding (SV2A^Y46A^). Unclustered Syt1 may be more accessible to the endocytic machinery, as the absence of SV2A potentiates dynamin recruitment and subsequent uptake of Syt1 into recycling SVs. Furthermore, the lack of bidirectional control of the retrieval of Syt1 and SV2A may be due to both molecules engaging different sets of endocytic adaptor molecules, which may lead to differing patterns of surface nanoclustering. For example, Syt1 has a unique interaction with stonin2, an endocytic adaptor that appears to exclusively shepherd Syt1 into clathrin-coated pits in concert with AP2 (Diril et al., 2006). Similarly to what we observed following SV2A knockdown, loss of stonin2 also accelerates the rate of Syt1 endocytosis (Kononenko et al., 2013).

### SV2A-mediated nanoclustering regulates the targeting of Syt1 into recycling vesicles

Several factors may account for the accelerated kinetics of unbound Syt1 in the absence of SV2A. One possible explanation for fast-paced internalisation of unclustered Syt1 into SVs may be a result of the increased recruitment of endocytic machinery to the PM. In support of this, our results demonstrate that the activity-dependent recruitment of dynamin to, and immobilization on, the PM is potentiated in the absence of SV2A. Another potential explanation is that the accelerated re-entry of Syt1 into recycling SVs may occur via ultrafast endocytosis – a mode of SV recruitment in which dynamin1 plays a critical role (Imoto et al., 2022; Watanabe et al., 2013). SV2A-Syt1 nanoclusters may therefore act as a sequestration hub, that rate-limits endocytosis by controlling the level of dynamin1 recruitment. As such, surface nanoclustering may act as a transition state for Syt1, allowing neurons to fine-tune endocytosis and titrate Syt1 internalisation back into recycling SVs. This sequestration process may facilitate titration due to steric limitations owing to the size of Syt1 nanoclusters, which are significantly larger than both clathrin-coated pits (0.065-0.125 µm) (Kirchhausen and Harrison, 1981) and recycling SVs (0.040 µm in diameter) (Zhang et al., 1998). Removing AP2 binding (SV2A^Y46A^) increased the interaction of SV2A with Syt1, raising the possibility that Syt1 interaction restricts SV2A’s access to the endocytic machinery. Endocytic events are also controlled by Syt1 (Li et al., 2017; McAdam *et al*., 2015; Nicholson-Tomishima and Ryan, 2004; Poskanzer et al., 2003), as the C2 domains of Syt1 regulate the kinetics of vesicle internalisation in a Ca^2+^-dependent manner (Yao et al., 2012). As the binding of SV2A to the C2B domain of Syt1 is negatively regulated by Syt1’s interaction with Ca^2+^ (Schivell et al., 2005), SV2A may compete with Ca^2+^ for C2B binding, thereby acting as a clamp. Therefore, it is tempting to speculate that the pace of Syt1-mediated endocytosis may be decreased through competitive interaction with Ca^2+^ and nanoclustering with SV2A.

The potentiated recruitment of dynamin1 to the PM upon dispersal of Syt1 from nanoclusters suggests that unclustered Syt1 associates with the endocytic machinery to speed up the rate of Syt1 internalisation. However, the presence of alternative trafficking routes, of differing endocytic kinetics, may also account for the fast-paced recruitment of Syt1 into recycling SVs. We demonstrate that SV2A-Syt1 interaction controls the intracellular sorting of Syt1 to Rab5-positive early or bulk endosomes, with clustered Syt1^WT^ preferentially associating with Rab5 following stimulation, in comparison to the SV2A binding mutant (Syt1^K326A/K328A^). Therefore, Syt1 nanoclustering likely restricts its access to smaller endocytic pits, instead redirecting it into Rab5-positive early endosomes (Wucherpfennig et al., 2003). This rerouting may play a physiological role in preventing the build-up of SV2A-Syt1 nanoclusters stranded on the PM during repetitive rounds of SV fusion. This mechanism likely favours activity-dependent internalisation of SV2A-Syt1 into bulk endocytic compartments that are positive for Rab5 (Kokotos et al., 2018) and large enough to engulf these nanoclusters. Bulk endosomes, have been shown to act as a sorting station for SV cargoes in which proteins can either be rerouted back to the reserve pool of SVs (Cheung et al., 2010) or trafficked to the endo-lysosomal system (Ivanova et al., 2021). Therefore, activity-dependent bulk endocytosis of SV2A-Syt1 during sustained neurotransmission may act as an intermediate step in the reformation of SVs.

### Functional consequences of SV2A-Syt1 nanoclustering on neurotransmission

The role of the SV2 family of proteins in neurotransmission is not well understood. Studies suggest that SV2A controls the size of the readily releasable pool of SVs (Custer et al., 2006), primes Ca^2+^-dependent SV fusion and regulates short-term synaptic plasticity (Chang and Südhof, 2009). Thus, as nanoclustering is theorised to determine the cellular function of proteins, our findings suggest that SV2A regulates Syt1 function by promoting its nanoclustering in both recycling vesicles and at the PM, ultimately controlling neurotransmitter release and plasticity. Understanding the molecular steps involved in this process provides insight into the role of SV2A and Syt1 during synaptic dysfunction, neurological disorders and potential avenues for therapeutic treatment. The critical importance of SV2A in vesicular targeting is described in a related study, which demonstrates that SV2 acts as a gateway for the entry of botulinum neurotoxin type-A (BoNT/A) into neurons, whereby BoNT/A hijacks SV2A-Syt1 nanoclusters to promote its own internalisation into SVs (Joensuu et al., 2022).

In summary, our results demonstrate that following exocytosis, PM-stranded vesicular proteins Syt1 and SV2A interact with each other to form nanoclusters independently of the endocytic machinery. These SV2A-Syt1 nanoclusters act upstream of endocytosis by rate-limiting the recruitment of the endocytic machinery, thereby controlling the rate of their retrieval into recycling SVs.

## Materials and methods

### Lead contact

Information and requests for reagents, materials and resources should be directed to and will be fulfilled by Professor Frédéric A. Meunier.

### Plasmids

Syt1-pH was a gift from Volker Haucke. Syt1^K326A/K328A^-pH, SV2A^WT^-pH, SV2A^T84A^-pH, SV2A-mCer, SV2A^T84A^-mCer and pSUPER vectors that co-express SV2A shRNA (shRNA sequence -GAATTGGCTCAGCAGTATG) with either mCer or Syt1-pH were described previously (Zhang et al., 2015). Rab5-mRFP was a gift from Giuseppe Balistreri. SV2A^Y46A^-pH and Syt1-HA were generated as reported in (Harper et al., 2020). mEos4b-Clathrin-15 was a gift from Michael Davidson (Addgene plasmid #57506; http://n2t.net/addgene:57506; RRID:Addgene 57506). SV2A-mEos2 cloning was carried out by the Protein Expression Facility, The University of Queensland (Brisbane, Australia). This involved replacing pHluorin from SV2A-pH with mEos2, which was inserted between amino acids 197 and 198. SV2A^Y46A^-mCer and SV2A^T84A/Y46A^-mCer were generated from either SV2A-mCer or SV2A^T84A^-mCer using the primers forward -GCATCCAGTGATGCT<u>G</u>CTGAGGGCCATGACGAG; Y46A reverse – CTCGTCATGGCCCTCAG<u>C</u>AGCATCACTGGATGC (mutated bases underlined). Site-directed mutagenesis was performed to introduce T112A into Syt1^WT^ -pH backbone using Quick-Change Lightning site directed mutagenesis kit (Aligent, #210518) as per manufacturer’s instructions. Primers used for Syt1 T112A Forward -5’ GAAAGACTTAGGGAAG<u>G</u>CCATGAAGGATCAGGC 3’ Reverse: 5’ GCCTGATCCTTCATGG<u>C</u>CTTCCCTAAGTCTTTC 3’. For Dyn1-mEos2, Dyn1 was amplified from pEGRF-N1-hDyn1 with the following primers: Forward -5’ TCGAATTCTGATGGGCAACCGCGGC 3’ and Reverse – 5’ GTGGATCCCGGGGGTCACTGATAG 3’. PCR products and pmEos2-N1 (Kasula *et al*., 2016) were digested with EcoRI (New England Bioscience, #R0101) and BamHI (New England Bioscience, #R0136), gel extracted and ligated with T4 ligase (New England Bioscience, #M0202) to make pmEos2-N1-hDyn1. The pEGRF-N1-hDyn1 plasmid was a generous gift from Phil Robinson. Positive clones were verified by sanger sequencing at the Australian Genome Resource Facility (AGRF, Brisbane).

### Animal maintenance

Experiments were performed using neurons derived from wild-type C57BL/6J mice (sourced from an in-house colony at the Queensland Brain Institute). For experiments in Brisbane, all work was carried out in accordance with the Australian Code and Practice for the Care and use of Animals for Scientific Purposes and approved by The University of Queensland Animal Ethics Committee (UQAEC -QBI/254/16/NHMRC). All C57BL/6J mice were housed with a 12 h light/dark cycle (light exposure between 7:00-19:00). Breeders were fed with autoclaved mouse and rat cubes (Specialty Feeds). All animal euthanasia was performed by approved standard operating procedures approved by the UQAEC. Adult mice were culled by cervical dislocation; embryos were euthanised by hypothermia and decapitation. For experiments in Edinburgh, animal work was performed in accordance with the UK Animal (Scientific Procedures) Act 1986, under Project and Personal Licence authority and was approved by the Animal Welfare and Ethical Review Body at the University of Edinburgh (Home Office project licence – 70/8878). All animals were killed by schedule 1 procedures in accordance with UK Home Office Guidelines; adults were killed by cervical dislocation followed by decapitation, whereas embryos were killed by decapitation followed by destruction of the brain. Wild-type C57BL/6J mice were sourced from an in-house colony at the University of Edinburgh. All mouse colonies were housed in standard open top caging on a 14 h light/dark cycle (light exposure between 07:00-21:00). Breeders were fed RM1 chow, whereas stock mice were maintained on RM3 chow.

### PC12 cell culture

Pheochromocytoma (PC12) cells were cultured in DMEM (Gibco, Life Technologies, #11995-065) containing 5 % heat-inactivated fetal bovine serum (FBS) (Bovogen, #SFBS-HI), 0.5 % GlutaMax supplement (Gibco, Life Technologies #35050061) and 5 % heat-inactivated horse serum (Gibco, Life Technologies #26050088) and maintained at 37°C with 5 % CO_2_. Transfections were performed using Lipofectamine LTX (Invitrogen, #94756) and Plus Reagent (Invitrogen, #10964-021). A day after transfection, cells were plated into glass-bottom culture dishes (Cellvis, CA, USA, #D29-20-1.5N) coated with poly-D-lysine (PDL, Sigma-Aldrich, #P1024-50MG).

### Hippocampal cell culture

Prior to dissection, 29 mm glass-bottom dishes (Cellvis, CA, USA, #D29-20-1.5N) were coated in poly-L-lysine (PLL, Sigma-Aldrich, #P2636-100MG) and left to incubate (37°C, 24 h). Hippocampal neurons were dissected from E16 embryos from C57BL/6J mice as previously described (Joensuu et al., 2017). Dissection was carried out in 1X Hank’s buffered salt solution, 10 mM HEPES pH 7.3 (Gibco, Life Technologies #15630-080), 100 U/ml penicillin-100 μg/ml streptomycin (Gibco, Life Technologies #15140-122). Digestion of hippocampal tissue was carried out using trypsin (0.25 % for 10 min, Sigma-Aldrich, #T4799-5G) and subsequently halted using FBS (5 %) with DNase I (Sigma-Aldrich, #D5025-375KU). Suspension was triturated and centrifuged (1500 rpm, 7 min), resuspended in plating medium (100 U/ml penicillin-100 μg/ml streptomycin, 1X GlutaMax supplement, 1X B27 (Gibco-Aldrich, Life Technologies, #17504-044) and 5 % FBS in neurobasal media (Gibco, Life Technologies #12348-017)). Seeding of neurons was carried out in glass-bottom dishes (1 × 10^5^ neurons covering the central glass bottom of each dish). Subsequently, plating media was fully replaced (2-4 h post-seeding) with culturing media (100 U/ml penicillin-100 μg/ml streptomycin, 1X GlutaMAX supplement, 1X B27 in neurobasal media). Hippocampal neurons 13-15 days-in-vitro (DIV) were transfected with plasmid (2 h, 2-3 µg per dish) with Lipofectamine 2000 (Invitrogen, #52887, minimum 24 h). For pH imaging experiments, dissociated primary hippocampal-enriched neuronal cultures were prepared from E16.5-18.5 embryos from wild-type C57BL/6J mice of both sexes as outlined (Harper et al., 2017; Zhang et al., 2015). In brief, isolated hippocampi were digested in 10 U/mL papain in Dulbecco’s phosphate-buffered saline (PBS), washed in Minimal Essential Medium (MEM) supplemented with 10 % FBS, and triturated to single cell suspension. This cell suspension was plated at 3-5 × 10^4^ cells on PDL and laminin-coated 25 mm coverslips. Cells were transfected on DIV 7-9 with Lipofectamine 2000 as per manufacturer’s instructions.

### Super-resolution microscopy

For live single particle tracking, neurons were placed in low K^+^ imaging buffer (5.6 mM KCl, 2.2 mM CaCl_2_, 145 mM NaCl, 5.6 mM D-Glucose, 0.5 mM MgCl_2_, 0.5 mM ascorbic acid (Sigma-Aldrich, #A5960), 0.1% bovine serum albumin (BSA), 15 mM HEPES, pH 7.4) at 37°C on a Roper Scientific Ring-TIRF microscope with a CPI Apo 100 x/1.49 N.A. oil-immersion objective (Nikon Instruments, NY, USA) with a Perfect Focus System (Nikon Instruments). Imaging was carried out using Evolve 512 Delta EMCCD cameras (Photometrics, AZ, USA), an iLas2 double laser illuminator (Roper Scientific, FL, USA), a quadruple beam splitter (ZT405/488/561/647rpc; Chroma Technology, VT, USA) and a QUAD emission filter (ZET405/488/561/640m; Chroma Technology). For imaging molecules on the plasma membrane, universal Point Accumulation Imaging in Nanoscale Topography (uPAINT) was carried out (Giannone et al., 2010). Neurons were stimulated with high K^+^ buffer (56 mM KCl, 0.5 mM MgCl_2_, 2.2 mM CaCl_2_, 95 mM NaCl, 5.6 mM D-Glucose, 0.5 mM ascorbic acid, 0.1 % BSA, 15 mM HEPES, pH 7.4) containing atto647N-labelled anti-GFP nanobodies (atto647N-NB) (Synaptic Systems, #N0301-AF647-L) at 3.19 pg µl^-1^. Treatments with Dyngo4a (Abcam, #ab120689) in DMSO were carried out at 30 μM in low K^+^ buffer for 30 min prior to addition of atto647N-NB in high K^+^ buffer. Control experiments involved a 30 min treatment of DMSO vehicle in low K^+^ buffer without Dyngo4a prior to application of atto647N-NB in high K^+^ buffer. For visualisation of both Syt1 nanoclusters and clathrin, uPAINT imaging of Syt1-pH-atto647N-NB was carried out in tandem with single particle tracking Photoactivated Localization Microscopy (sptPALM) of clathrin-mEos4b. The mEos4b fluorophore was excited through the application of a 405 nm laser, which triggered its photoconversion from green to red. Photoconverted mEos4b was simultaneously imaged using excitation with a 561 nm laser.

For imaging of Syt1-pH-atto647N-NB internalised in recycling vesicles, sub-diffractional Tracking of Internalised Molecules (sdTIM) was used. Neurons were pulsed for 5 min in high K^+^ buffer containing atto647N-NB (3.19 pg µl^-1^), before being washed with low K^+^ imaging buffer (5x) and left under resting conditions for a further 5 min. Subsequently, neurons were incubated in imaging buffer containing active TEV protease (Invitrogen, 12575-015) (60U, 15 min). For 3D-Structured Illumination Microscopy (3D-SIM), neurons underwent sdTIM imaging and were subsequently fixed in paraformaldehyde (PFA, ProSciTech, #C004) (4% in PBS) (15 min), washed in PBS (5x) and mounted in non-hardening antifade mounting medium (Vectashield, H-1000). 3D-SIM acquisitions were taken on an Elyra PS.1 microscope (100X objective). A series of z-stacks (23 slices, 0.1 µm intervals, 1024×1024 pixels) were taken sequentially in three channels (488, 561 and 640 nm). For channel alignment (xyz), z-stacks were taken of multifluorescent Tetraspeck beads (Invitrogen, T7279).

### Immunocytochemistry

For immunolabelling of endogenous SV2A, transfected neurons were fixed in PFA (4 %) in PBS (20 min). Neurons were washed in PBS (3x) and incubated in blocking buffer consisting of BSA (1% in PBS) for 30 min. Neurons were subsequently incubated with a rabbit anti-SV2A antibody (Abcam, ab32942) (1:200, 1 h) in blocking buffer. Neurons underwent further washes in PBS (3x) and were labelled with anti-rabbit alexa594 (Invitrogen, A11035) (1:1000, 30 min), prior to undergoing additional PBS wash steps (3x). Fixation, blocking, antibody incubation and wash steps were all carried out at room temperature.

### Fluorescence imaging

Timelapse recordings of pHluorin (pH) were performed as previously described (Harper et al., 2020). Hippocampal neurons were transfected with pH- and mCer-tagged proteins of interest at DIV 7. Between DIV 13-15, neurons were mounted in a Warner Instruments (Hamden, CT, USA) imaging chamber on a Zeiss Axio Observer D1 or Z1/7 inverted epifluorescence microscope (Cambridge, UK) with a Zeiss EC Plan Neofluar 40x/1.30 oil immersion objective fitted with an AxioCam 506 mono camera (Zeiss). The pH fluorophore was excited at 500 nm. SV2A-mCer was excited at 430 nm. Visualisation of mCer and pH was carried out using a long-pass emission filter (>520 nm). Field stimulation was performed on neuronal cultures using a train of 300 action potentials (APs) at 10 Hz (100 Ma, 1 ms pulse width). Images of pH were taken at 4 s intervals. After 180 s post-stimulation, neurons were perfused with an alkaline buffer containing NH_4_Cl (50 mM), increasing the intracellular pH thereby revealing the fluorescence levels of the total pH-tagged SV2A population. For TIRF imaging of Dyn1-GFP, PC12 cells were imaged three days following transfection incubated in buffer A (145 mM NaCl, 5 mM KCl, 1.2 mM Na_2_HPO_4_, 10 mM D-glucose and 20 mM HEPES, pH 7.4). Subsequently, transfected cells were imaged (37 °C, 5 % CO_2_) using the inverted Roper Scientific TIRF microscope (Roper Scientific). TIRF imaging of Dyn1-GFP was performed with a 491 nm laser at 50 Hz (16,000 frames) using MetaMorph 7.7.8 (Molecular Devices, CA, USA).

### Co-immunoprecipitation and western blotting

HEK-293T cells were maintained at 37 °C, 5 % CO_2_ in culture media (MEM) (Invitrogen, 41966-029), 10 % FBS, 1 % penicillin/streptomycin). Cells were co-transfected with SV2A-mCer (SV2A^WT^-mCer, SV2A^T84A^-mCer or SV2A^Y46A^-mCer) and Syt1-HA using Lipofectamine™ 2000 (48 h) (ThermoFisher # 11668019, as per manufacturer’s instructions). Cells were solubilised in HEPES buffer (50 mM HEPES pH 7.5, 0.5 % Triton X-100, 150 mM NaCl, 1 mM EDTA, 1 mM PMSF, protease inhibitor cocktail) for 1 h prior to centrifugation (17,000 g for 10 min), from which the resulting supernatant was isolated and treated with GFP-Trap beads (Chromotek, Germany) and rotated at 4 °C for 2 h, followed by additional wash (3X) in HEPES buffer. Samples were incubated in SDS sample buffer (10 min at 65 °C) and loaded on an SDS-PAGE gel for western blotting, which was carried out in accordance with previous studies (Anggono et al., 2006). Primary antibodies used were anti-GFP rabbit (Abcam, ab6556, 1:4000) and anti-HA rabbit (ICLlab, RHGT-45A-Z, 1:20000). IRDye secondary antibody (800CW anti-rabbit IgG (#925-32213, 1:10000) and Odyssey blocking PBS buffer (#92740000) were from LI-COR Biosciences (Lincoln, Nebraska, USA). Blots were visualised using a LI-COR Odyssey fluorescent imaging system (LI-COR Biotechnology, Cambridge, UK). Band densities were determined using LI-COR Image Studio Lite software (version 5.2). The amount of Syt1-HA co-immunoprecipitated was normalised to the amount of input protein. These values were then normalised to the amount of immunoprecipitated SV2A-mCer.

### Image processing

For single particle tracking, image processing was carried out in PALMTracer, a custom-written software that operates in MetaMorph 7.7.8 (Molecular Devices, CA, USA) (Kechkar et al., 2013). Regions of interests (ROIs) were drawn around nerve terminals defined as hotspots of increased pHluorin fluorescence. As exocytic fusion occurs exclusively at the level of the synapse, we have systematically used pHluorin unquenching to delineate the presynapse. Trajectories lasting a minimum of eight frames were selected and reconstructed. The mean square displacement (MSD) was calculated by fitting the equation MSD(*t*)=*a*+4*Dt* (where *D* is diffusion coefficient, *a* is y intercept and *t* is time), with MSD quantified over a 200 ms period. The diffusion coefficient was calculated and divided into mobile and immobile populations with a diffusion coefficient of log_10_ > -1.45 µm^2^ s^-1^ considered as mobile (Constals et al., 2015; Joensuu *et al*., 2016). A SharpViSu tool was used to perform drift correction for cluster analysis (Andronov et al., 2016). A custom-built python tool was used to perform NAnoscale SpatioTemporal Indexing Clustering (NASTIC) on our track files to determine the size, density, and apparent lifetime of Syt1 and SV2A nanoclusters. NASTIC generates a series of overlapping, spatiotemporal bounding boxes around trajectories used to determine cluster formation (Wallis et al., 2021). Nanoclusters were thresholded at a radius of 0.15 µm, with anything greater excluded from analysis.

Offline data processing of pH-transfected neurons was performed using Fiji Is Just ImageJ (Fiji) software (Schindelin et al., 2012). A script based on background thresholding was used to select nerve terminals, which placed ROIs of identical size over those responding to stimulation. Average fluorescent intensity was measured over time using the Time Series Analyzer plugin before screening ROIs using a customised Java program that allows for visualisation of the fluorescent responses and removal of aberrant traces from the data. Subsequent data analyses were performed using Microsoft Excel, Matlab (Cambridge, UK) and GraphPad Prism 6.0 (La Jolla, CA, USA) software. The change in activity-dependent pHluorin fluorescence was calculated as F/F_0_ and normalised to the peak of stimulation.

3D-SIM processing and channel alignment were performed using Zen 2012 SP2 Black (version 11.0, ZEISS). Rab5-mRFP clusters were observed along the axon of each neuron. Colocalization of points and surfaces was carried out in IMARIS (version 9.6.0). A series of 3D surfaces were defined for Rab5 cluster points (561 nm) across each z-stack based on the signal intensity across the neuron, with neuronal morphology defined based on Syt1-pH signal. For each surface, the points corresponding to internalised Syt1-pH-atto647N-NB molecules (642 nm) were identified.

### Statistics and data analysis

A Students *t*-test was performed for comparison between two groups (Fig. 1-2, Fig. 3A, Fig. 3C iv-vi, D i-iv, Fig. 4C i-iii, Fig. 4D iv, Fig. 6-7). A one-way ANOVA was performed followed by a post-hoc Tukey’s test for multiple comparison data with a gaussian distribution of residuals (Fig. 3C i-iii, Fig. 4B, Fig. 4D ii, Fig. 5B). For non-parametric analysis assuming no gaussian distribution, a Kruskal-Wallis test with Dunn’s test for multiple comparisons was carried out (Fig. 5C-D). The level of significance was set to p<0.05. Error bars represent standard error of the mean (SEM). All n values correspond to independent cells/neurons unless specified (e.g., Fig. 4D i-iii where n corresponds to individual nanoclusters).

## Data and code availability

A custom-built Python tool for NASTIC analysis was can be found at (Wallis et al., 2021) with the Python code available at https://github.com/tristanwallis/smlm_clustering. Analysis of pHluorin fluorescence in nerve terminals was carried out using a custom-made script based on background thresholding. This was used to select nerve terminals, which placed regions of interest of identical size over those responding to stimulation (Harper et al., 2020).

## Author contributions

C.S., M.A.C. and F.A.M. conceived and designed the project. The manuscript was written by C.S. and F.A.M. Additional manuscript edits were provided by M.A.C., M.J. and T.P.W.. Dynamin recruitment assays in PC12 cells were performed by A.J.. Single-particle tracking (uPAINT, sdTIM and sptPALM), 3D-SIM imaging and processing was carried out by C.S. with the help of and N.Y.. Site-directed mutagenesis of Syt1T112A-pH was carried out by R.S.G.. Fluorescence recordings of Syt1-pH and SV2A-pH were performed by C.H. and C.K., while E.C.D. performed the co-immunoprecipitations. T.W. designed the cluster analysis software. IMARIS colocalization was performed by A.M.. Technical assistance was provided by R.M.M. and N.Y. Drift correction was performed by M.J..

## Acknowledgments

The super-resolution imaging was carried out at the Queensland Brain Institute’s (QBI’s) Advanced Microimaging and Analysis Facility. We thank Dr Adekunle Bademosi, Dr Rumelo Amor and Dr Arnaud Gaudin for technical assistance with SIM imaging and analysis. Many thanks to Dr Nick Valmas for providing the illustrations of Syt1-pH (Fig. 1) and SV2A-pH (Fig. 4), and Dr Alex McCann for assistance with figures and for editorial revisions. Thanks to Ms Barbara Duda for editorial revisions. We thank Dr. Merce Salla-Martret for technical assistance with Dyn1-mEos2 cloning. We thank the Clem Jones Centre for Ageing Dementia Research (CJCADR) for its support. Thanks to Professor Phil Robinson for providing the Dyngo4A used in this study. This work was also supported by an Australian Research Council Discovery Project grant (170100125), an Australian Research Council Linkage Infrastructure, Equipment, and Facilities grant (LE130100078), a National Health and Medical Research Council (NHMRC) grant (1139316) to F.A.M. A grant from the Biotechnology and Biological Sciences Research Council (BB/L019329/1) to M.A.C. Research was funded in part by the Wellcome Trust [Investigator Award to M.A.C. (204954/Z/16/Z)]. For the purpose of open access, the author has applied a CC-BY public copyright license to any author accepted manuscript version arising from this submission. M.J. is supported by an Australian Research Council Discovery Early Career Researcher Award (DE190100565). R.M.M. was supported by the Clem and Jones Foundation, the State Government of Queensland, and the NHMRC Boosting Dementia Research Initiative. C.S. was supported by the Research Training Program (RTP) Scholarship and a QBI top-up scholarship. F.A.M. is an NHMRC Senior Research Fellow (1155794). We thank Dr Rona Wilson and Mr Hamish Runicman for assistance with neuronal culture preparation and Dr Donal Stewart for the development of a Java program to visualise and screen data. The authors of declare no conflict of interest.

---

**Supplementary figure 1.**
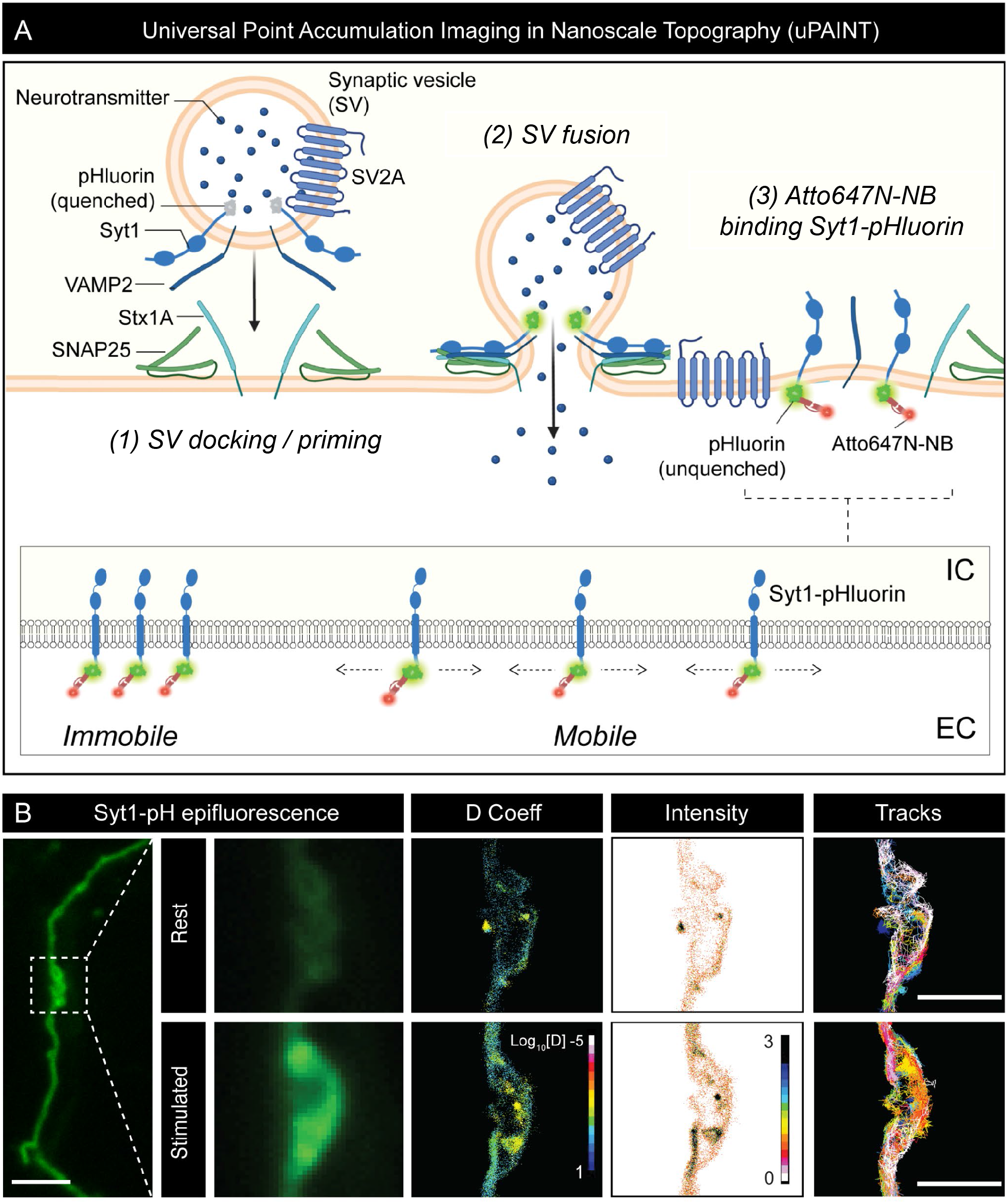
(Figure 1). Universal Point Accumulation Imaging in Nanoscale Topography (uPAINT) methodology. **(A)** (1) SV docking and priming: SNARE proteins assemble. While in the acidic environment of SVs, the pHluorin tag of Syt1-pH is quenched. (2) SV fusion (exocytosis): the SV fuses with the PM causing neurotransmitter release. Syt1-pH becomes unquenched upon exposure to the neutral extracellular environment, indicative of SV fusion. (3) Atto647N-NB binding to Syt-pH: following fusion, Syt1-pH is on the PM and accessible to anti-GFP atto647N-NB. This enables uPAINT imaging of Syt1-pH-atto647N-NB on the PM, which transitions between an unclustered mobile state and a clustered immobile state. **(B)** uPAINT imaging of hippocampal axons and nerve terminals pre- and post-stimulation (high K^+^). Increased Syt1-pH fluorescence can be observed following stimulation due to the post-exocytic unquenching of the pHluorin tag. Corresponding diffusion coefficient (D Coeff), intensity and trajectory maps of Syt1-pH-atto647N-NB are shown. For the D Coeff panels, the colour bar represents log_10_[D] (µm^2^s^-1^), with warmer colours indicating points of low mobility. Scale bar = 2 µm (axon) and 1 µm (presynapse).

**Supplementary figure 2.**
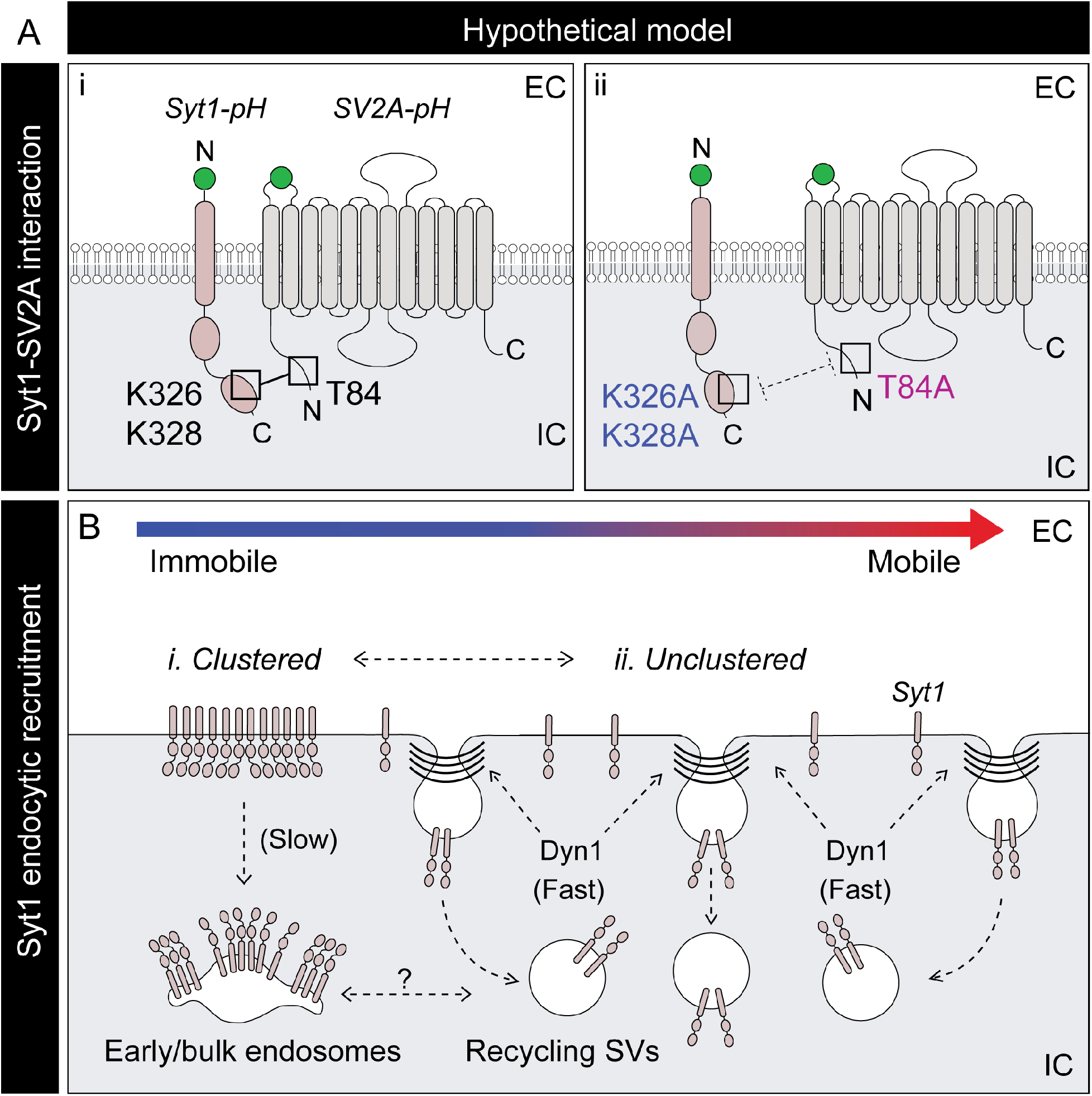
(Figure 7). Hypothetical model: SV2A-bound Syt1 nanoclusters control the selective targeting of Syt1 to recycling synaptic vesicles. **(A)** Schematic showing (i) the SV2A-Syt1 interaction that forms between the K326/K328 epitopes at the polybasic region of the C2B domain of Syt1^WT^-pH and the T84 epitope at the N-terminal tail of SV2A^WT^-pH, and (ii) disruption of the SV2A-Syt1 interaction via mutation (Syt1^K326A/K328A^ or SV2A^T84A^). **(B)** Schematic depicting the effect of the SV2A-Syt1 interaction on the nanoclustering and subsequent internalisation of Syt1. (i) Syt1 forms nanoclusters on the PM that are highly immobile and may be preferentially sorted into Rab5-positive early/bulk endosomes. (ii) Unclustered Syt1 is more mobile and can recruit dynamin1 to promote its internalisation into recycling SVs, which occurs at a faster rate.

